# Multimodal HLA-I genotype regulation by human cytomegalovirus US10 and resulting surface patterning

**DOI:** 10.1101/2023.01.10.523457

**Authors:** Carolin Gerke, Liane Bauersfeld, Valerie Oberhardt, Christopher Sebastian Jürges, Robbert M. Spaapen, Claudio Mussolino, Vu Thuy Khanh Le-Trilling, Mirko Trilling, Lars Dölken, Florian Erhard, Maike Hofmann, Hartmut Hengel, Frank Momburg, Anne Halenius

## Abstract

To control human cytomegalovirus (HCMV) infection, NK cells and CD8^+^ T-cells are crucial. HLA class I (HLA-I) molecules play a central role for both NK and T-cell responses and are targets of multifaceted HCMV-encoded immunoevasins. A so far insufficiently studied HLA-I immunoevasin is the glycoprotein US10. It was shown that US10 targets HLA-G, but it is unknown whether US10 contributes also to escape from classical HLA-I antigen presentation. Our biochemical analysis revealed that early during maturation, all investigated HLA-I (HLA-A/B/C/E/G) heavy chains are recognized and bound by US10. Remarkably, the consequences of this initial binding strongly depended on both the HLA-I geno- and allotypes: i) HLA-A molecules escaped down-regulation by US10, ii) tapasin-dependent HLA-B molecules exhibited impaired recruitment to the peptide loading complex and maturation, iii) HLA-C and HLA-G, but not HLA-A/B/E, strongly bound US10 also in their β_2_m-assembled form. Thus, US10 senses geno- and allotypic differences in a so far unparalleled and multimodal manner, suggestive of adaptation to HLA-I genotype differences. At a further level of complexity, in HCMV-infected fibroblasts inhibition of overlapping *US10* and *US11* transcription revealed an additional HLA-I specificity, suggesting targeting of HLA-I in a synergistically arranged manner. Our study unveils the exceptional HLA-I selectivity of HCMV-encoded US10 and underlines its contribution to immune escape.

## Introduction

The human cytomegalovirus (HCMV) belongs to the β-herpesviruses and establishes a life-long persistent infection in its host with alternating phases of latency and reactivation. Although clinical manifestations are mainly observed in immunocompromised patients, HCMV also affects the immune system of healthy individuals (Brodin et al., 2015). Expansion of memory-like NK cells as well as of CD8^+^ memory T cells are frequent observations in HCMV-positive individuals (Lubke et al., 2020; Rolle and Brodin, 2016; Waller et al., 2008). These cytotoxic lymphocytes are crucial for HCMV control (Sylwester et al., 2005; Venema et al., 1994). While specific antigenic peptide ligands presented by MHC class I (MHC-I) molecules on infected cells activate CD8^+^ T-cells, NK cells express various inhibiting and activating receptors that recognize MHC-I both in peptide-independent and - dependent manners.

In humans, MHC-I molecules are encoded by three classical (A, B, C) and three non-classical (E, F, G) human leucocyte antigen class I (HLA-I) loci located on chromosome 6 (Koller et al., 1989). Classical HLA-I are characterized by a high degree of polymorphism, whereas non-classical HLA-I show low levels of heterogeneity. The MHC-I maturation process begins with a co-translational translocation of the heavy chain (HC) into the endoplasmic reticulum (ER), where it folds and assembles with β_2_-microglobulin (β_2_m). To acquire a peptide ligand, MHC-I is assisted by the chaperones tapasin, ERp57 and calreticulin as well as the transporter associated with antigen processing (TAP), together forming the peptide loading complex (PLC) (Blees et al., 2017; Hulpke and Tampe, 2013). While TAP transports cytosolic peptides generated by the proteasome into the ER (Michalek et al., 1993), tapasin serves as an adapter chaperone linking TAP and HLA-I. Additionally, tapasin fulfils a crucial role in peptide editing and optimization (Wearsch and Cresswell, 2007; Williams et al., 2002). Peptide-loaded MHC-I is transported to the cell surface via the secretory pathway enabling immune cell surveillance of MHC-I expression and antigen presentation.

Different from most viruses, cytomegaloviruses encode several proteins targeting MHC-I molecules. For this, HCMV applies various strategies (Halenius et al., 2015): US2 and US11 initiate degradation of the MHC-I HC (Jones and Sun, 1997; Wiertz et al., 1996), US3 strongly retains MHC-I in the ER and blocks tapasin function (Jones et al., 1996; Park et al., 2004) and US6 blocks the peptide transport by TAP (Ahn et al., 1997; Hengel et al., 1997; Lehner et al., 1997). These immunoevasins belong to the *US6* gene family, also including US9, which targets the MHC-I-like molecule MICA*008 with its signal peptide (Seidel et al., 2021), and US7 and US8, which antagonize Toll-like receptor signaling (Park et al., 2019). Like all members of the *US6* gene family, US10 is a type I transmembrane glycoprotein and localized in the ER (Huber et al., 2002). Immunoprecipitation experiments demonstrated binding to MHC-I as well as a delay of MHC-I maturation in the presence of US10 (Furman et al., 2002). Furthermore, US10 was shown to be involved in the degradation of the non-classical MHC-I molecule HLA-G (Park et al., 2010). Altogether, a role of US10 as an MHC-I immunoevasin is clearly indicated, but it is incompletely understood to which extent US10 binding to HLA-I fulfils biological functions.

We previously decoded pronounced genotype-specific differences in HLA-I regulation by US11: HLA-A is strongly targeted by US11-mediated proteasomal degradation, whereas HLA-B can escape proteasomal targeting (Zimmermann et al., 2019). Thus, we hypothesized that also US10 possesses HLA-I genotypic, or even allotypic, preferences and aimed at elucidating the targeting spectrum of US10 to gain insights into its role in control of HLA-I functions.

Here, we describe a great US10 selectivity for HLA-I targets highlighting an intricate role for US10 as an HLA-I immunoevasin.

## Results

### Geno- and allotype-specific downregulation of HLA-I cell surface expression by US10

To gain deeper insight into the immunoevasive role of US10, we set out to determine its HLA-I preferences using a comprehensive and quantitative assay based on flow cytometry (Zimmermann et al., 2019). A panel of plasmids encoding N-terminally HA-tagged HLA-I molecules, or CD99 as control, were co-transfected with an EGFP-expressing vector encoding US10 or a control protein. HLA-I surface expression was determined by anti-HA staining of EGFP-positive cells at 20 hours post-transfection. The HLA-I-related molecules MICA*004, MR1, and the murine H-2K^b^ were not downregulated by US10, while US10 displayed a broad spectrum of HLA-I regulation (Fig. 1A-B). Intriguingly, all five tested HLA-A allotypes were resistant to US10-mediated downregulation. As expected, HLA-G surface expression was almost completely lost. Intriguingly, also the expression of other HLA-I molecules was greatly reduced, i.e. HLA-B*44:02, -C*05:01, and E, and to a slightly lesser extent HLA-C*04:01, -C*12:03, -B*08:01, and -B*15:03.

**Fig. 1.**
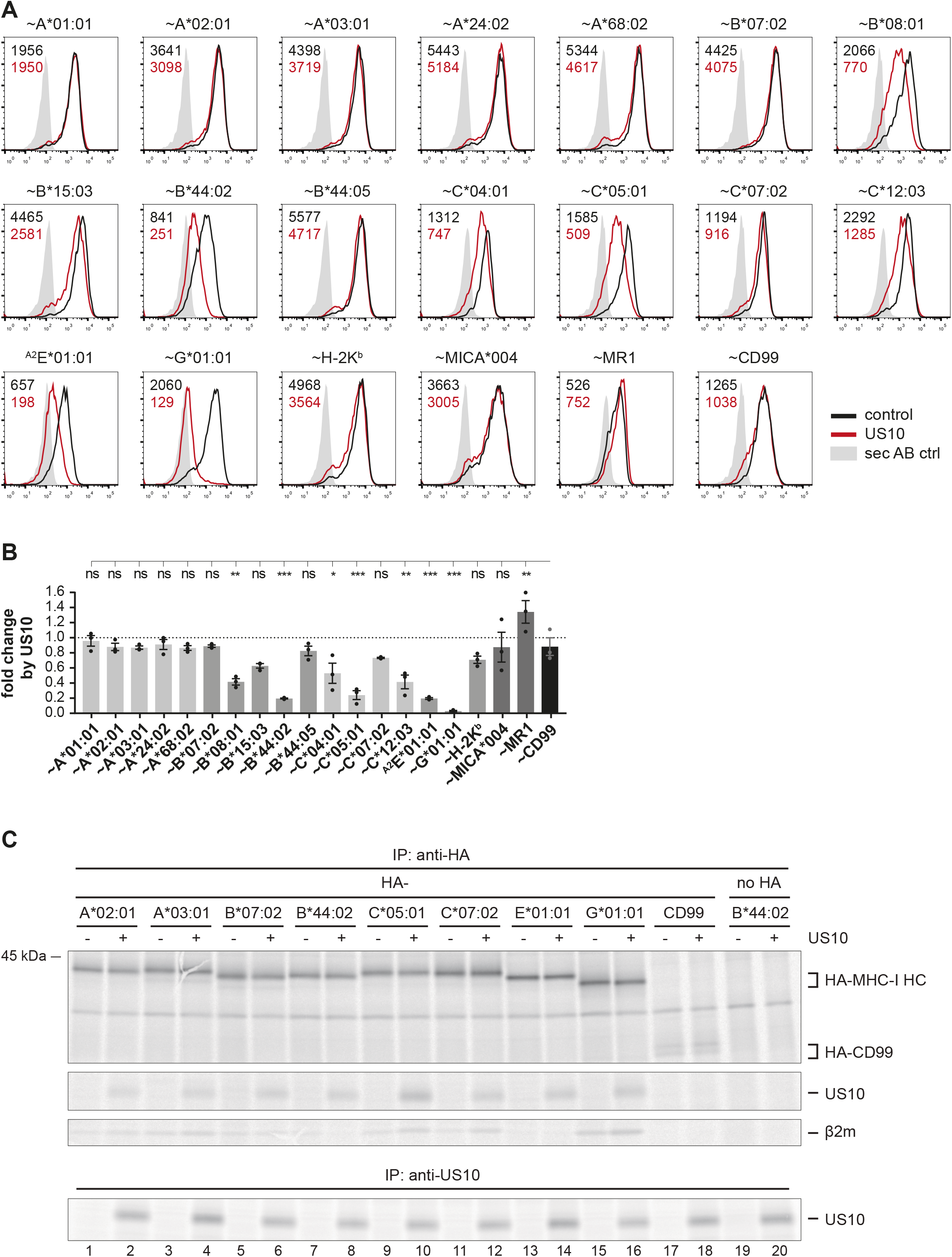
Geno- and allotype-specific regulation of HLA-I by US10. (A) HeLa cells were transiently co-transfected with plasmids encoding HA-tagged (∼) molecules or untagged HLA-^A2^E*01:01 (HLA-E was expressed with an HLA-A*02 signal peptide, a natural HLA-E ligand) and plasmids with either US10 or a control protein together with an IRES-*EGFP* cassette. Surface expression was determined by flow cytometry (anti-HA or anti-HLA-E) on EGFP-positive cells. Representative histograms are shown. (B) Fold change of surface expression by US10 was calculated as the ratio of the MFI of US10 expressing cells compared to control transfected cells. Dots represent individual values and bars mean values ± SEM from three independent experiments (biological replicates). Significance compared to the HA-CD99 control was calculated using one-way ANOVA followed by Dunnett’s multiple comparison test. (C) HeLa cells were transiently transfected as described in (A) and metabolically labeled for 2h. Immunoprecipitations using anti-HA or anti-US10 were performed and separated by SDS-PAGE with subsequent detection by autoradiography. One of two independent experiments (biological replicates) is shown.

To better understand the molecular basis for US10 targeting of HLA-I, we next analyzed whether the distinct effects observed on HLA-I allotypes was a result of variable US10 binding to HLA-I. To this end, US10 and HA-tagged HLA-I were transiently expressed and a co-immunoprecipitation assay with an anti-HA antibody was performed after metabolic protein labeling (Fig. 1C). HA-CD99 and untagged HLA-B*44:02 were included as negative controls. With comparable intensity, US10 bound to all tested HLA-I molecules (including allotypes of HLA-A, -B, -C, -E and -G) except for HLA-C*05:01, which co-immunoprecipitated US10 stronger than the other HLA-I allotypes.

Furthermore, in contrast to a previous observation (Park et al., 2010), we did not observe enhanced degradation of HLA-G in US10-expressing cells (Fig. 1C). This was also not the case for any other HLA-I molecule studied in this experiment. Thus, US10 binding to HLA-I did not explain the differences observed in HLA-I cell surface expression, nor did we observed destabilization of HLA-I by US10.

### US10 blocks HLA-I interaction with the PLC

It has previously been shown that US10 decreases the rate of HLA-I export from the ER, however it was not determined whether allomorph differences exist (Furman et al., 2002). Therefore, we assessed whether this effect is HLA-I selective. HeLa cells endogenously express HLA-A*68:02, - B*15:03, and -C*12:03. When separated by SDS-PAGE, HLA-A*68:02 can be distinguished from the two other HLA-I allotypes due to faster migration (Zimmermann et al., 2019). In HeLa cells stably expressing US10, we found that HLA-A*68:02 was clearly less retained in the ER/cis-Golgi by US10 than the other HLA-I as judged from acquisition of endoglycosidase H (EndoH, EH) resistant glycans in pulse-chase experiments (Fig. 2A-B).

**Fig. 2.**
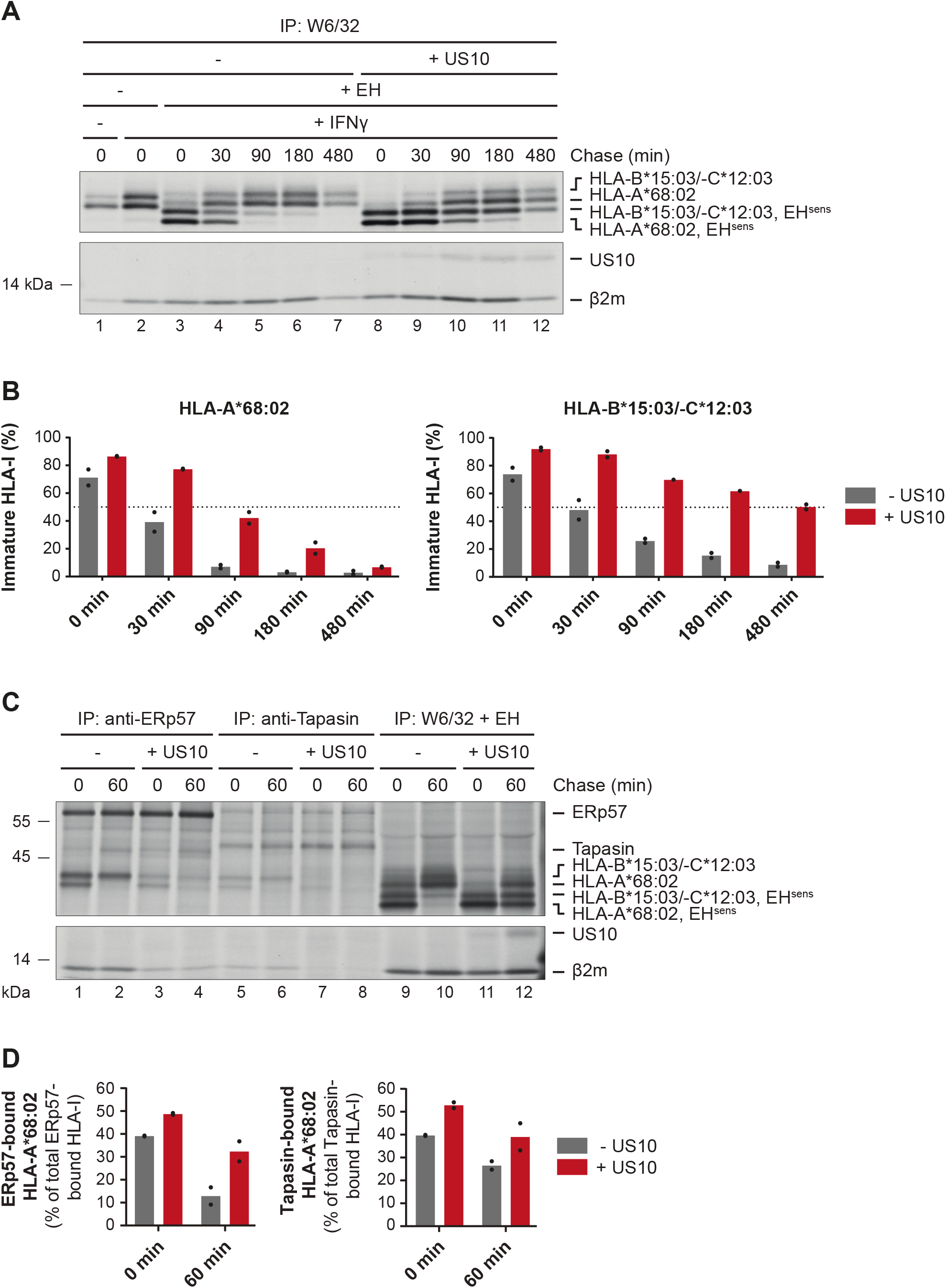
US10 blocks HLA-I interaction with the PLC. (A) Control HeLa cells or cells stably expressing US10 were induced by IFNγ overnight and subsequently metabolically labeled for 30 min and chased as indicated. After immunoprecipitation by W6/32, proteins were digested by EndoH (EH^sens^, EndoH-sensitive molecules) as indicated and separated by SDS-PAGE. Labeled proteins were detected by autoradiography. (B) The intensities of single HLA-I HC bands in (A) were quantified and the percentage of immature molecules as compared to the total amount (sum of immature and mature) was calculated and depicted from two independent experiments (biological replicates). (C) Immunoprecipitation from HeLa cells or cells stably expressing US10 was performed as in (A) but with modified chase times and without IFNγ treatment. Antibodies applied for immunoprecipitations are indicated. (D) Band intensities of HLA-I HCs in anti-ERp57 and anti-tapasin immunoprecipitations from (C) were quantified and the amount of the HLA-A*68:02 HC was calculated as the percentage of total PLC-bound HLA-I (sum of both HC bands). Dots represent individual values from two independent experiments (biological replicates).

A rate determining factor for the ER exit of MHC-I molecules is the average time required to load a stabilizing peptide in the binding groove. The dependency on the PLC for efficient peptide loading varies greatly among MHC-I allotypes. To assess whether the PLC is involved in the delay of MHC-I maturation caused by US10, HLA-I interaction with the PLC was analyzed in pulse-chase experiments (Fig. 2C). Co-immunoprecipitations with antibodies directed against ERp57 and tapasin revealed that the overall level of PLC-associated HLA-I was greatly reduced in cells expressing US10, despite the increased EndoH sensitivity of HLA-I in the presence of US10 (Fig. 2C, lanes 9-12), indicating that larger quantities of HLA-I molecules remained in the ER/cis-Golgi. This shows that US10 strongly interferes with HLA-I/PLC interaction. Interestingly, we observed that HLA-A*68:02 was less affected by this than HLA-B*15:03/C*12:03; the percentage of HLA-A*68:02 in the PLC as compared to the total amount of HLA-I in the PLC was increased in US10-expressing cells (Fig. 2D). Hence, HLA-A*68:02 access to the PLC is less disturbed by US10 than observed for HLA-B*15:03/-C*12:03.

Since US10 was not co-immunoprecipitated with the PLC, neither after a short (30 min, Fig. 2C) nor a long (2h, Suppl. Fig. 1) metabolic labeling, US10-bound HLA-I is likely excluded from the PLC. This is different from the other HCMV-encoded MHC-I immunoevasins since US3 and US11 both physically interact with the PLC (Park et al., 2004; Zimmermann et al., 2019). Furthermore, we demonstrate here that under the same experimental conditions US2 and US3 were co-immunoprecipitated by anti-ERp57, whereas this was not the case for US10 (Suppl. Fig. 1). Thus, US10 intervenes with MHC-I biogenesis at a unique level.

### The strength of HLA-B regulation by US10 correlates with HLA-B tapasin dependency

The strong US10 sensitivity of HLA-B*44:02 and -B*08:01, but not of HLA-B*44:05 and -B*07:02, and the finding that US10 blocked HLA-I interaction with the PLC, prompted us to analyze whether the level of HLA-I tapasin dependency correlated with US10 sensitivity. We engineered tapasin knock-out HeLa cells (Suppl. Fig. 2A) and compared the effect of tapasin deficiency with the effect of ectopic US10 expression (Fig. 3A and Suppl. Fig. 2B). To enhance TAP expression and peptide translocation to the ER in the absence of tapasin, cells were treated with IFNγ. The level of HLA-I tapasin dependency correlated well with data from others (Bashirova et al., 2020) and IFNγ treatment did not change the pattern of allotype-dependent regulation by US10. Consistent with the observation that HLA-A allotypes were resistant to US10, no correlation between tapasin dependency and US10 sensitivity was observed for HLA-A (Fig. 3B). In contrast, HLA-B tapasin dependency correlated strongly with US10-mediated downregulation. Curiously, even though HLA-C molecules were regulated in an allotype-dependent manner by US10, there was no correlation with tapasin dependency. This implied that US10 targeting of HLA-I allotypes is rather governed by their genotypes than by tapasin function. Moreover, the analysis of US10 function in tapasin knock-out cells showed that US10 was able to further reduce surface expression of US10 sensitive HLA-I such as HLA-B*15:03 and HLA-G*01:01 (Fig. 3C), indicating that the presence of tapasin is not a prerequisite for the function of US10.

**Fig. 3.**
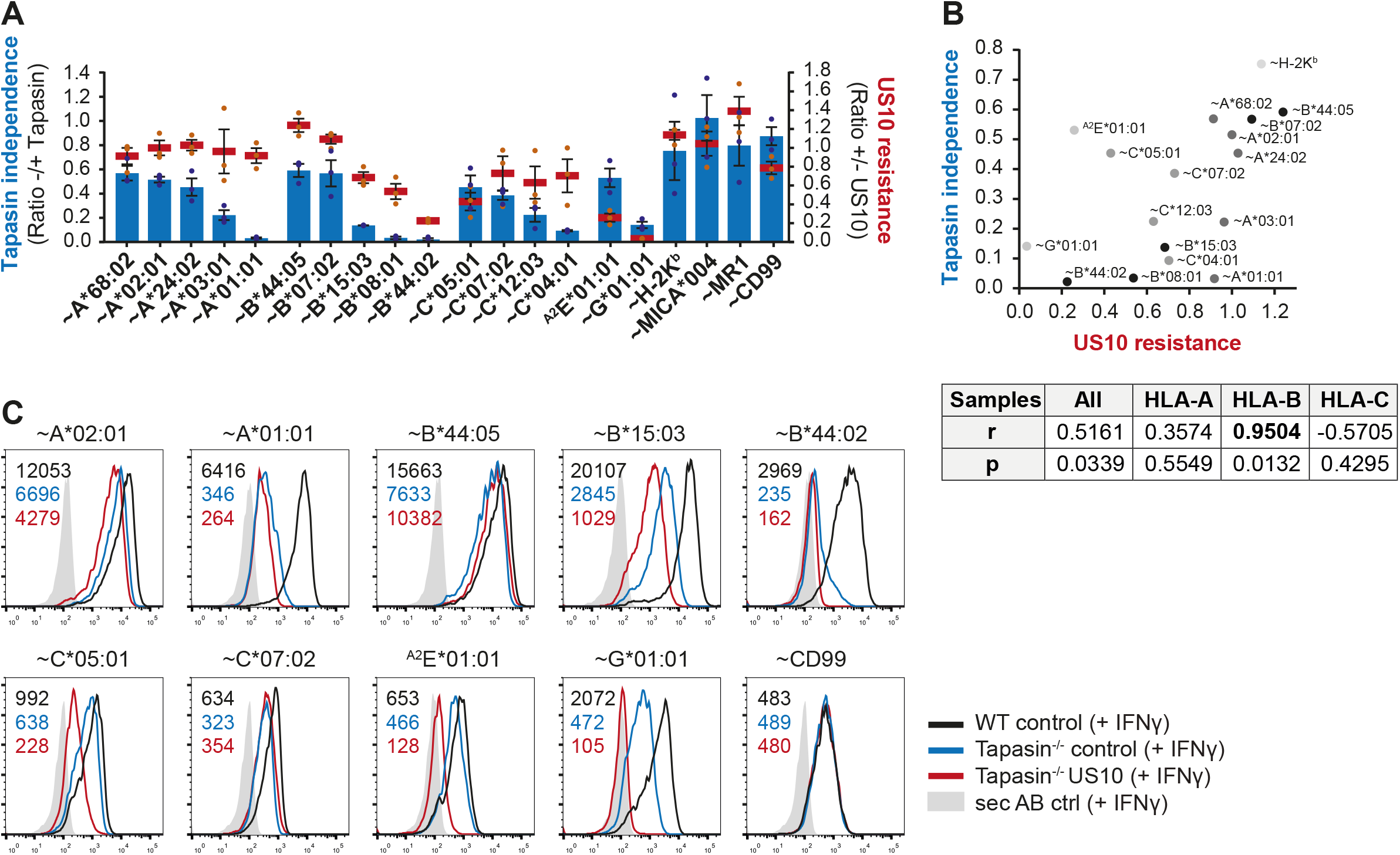
US10 control of HLA-B alloforms correlates with tapasin dependency. (A) Wild-type or tapasin knock-out HeLa cells were transiently co-transfected as indicated and treated with IFNγ overnight. Cell surface expression of the HA-tagged molecules or untagged HLA-^A2^E*01:01 was determined (representative histograms in Suppl. Fig. 2B) as in Fig. 1A. US10 resistance was calculated as the ratio of the MFI of US10-expressing cells as compared to control cells (red lines). Tapasin independence was calculated as the ratio of the MFI of tapasin knock-out compared to wild-type cells (blue bars). Dots represent individual values and bars mean values ± SEM from three independent experiments (biological replicates). (B) Two-tailed correlation analysis of the results from (A). (C) Flow cytometry analysis performed as in (A) including US10 in tapasin knock-out cells. Representative histograms from one of three independent experiments (biological replicates) are shown.

### US10 binding to HLA-I comprises two recognition modalities

The anti-HA antibody applied in the US10 binding analysis in Figure 1C binds to the HA-tagged HLA-I heavy chain independently of the maturation stage and does not reveal which conformational form of HLA-I is recognized by US10. We wondered whether conformation-dependent interactions could explain the HLA-I selective effects of US10. To address this question, we deleted endogenous HLA-I from HeLa cells using a lentiviral CRISPR/Cas9 system with a gRNA targeting a common sequence present in the HLA-I alleles expressed by HeLa cells (Supp. Fig. 3A-B). We then mutated the gRNA target site in HA-tagged HLA-I constructs and transiently co-expressed them in HeLa HLA-I knock-out cells together with US10 or a control protein. The transfected cells were metabolically labeled and binding of US10 was assessed with both anti-HA and the mAb W6/32, which recognizes only β_2_m-assembled HLA-I. Remarkably, US10 bound much stronger to β_2_m-assembled HLA-C and -G than to HLA-A, -B, and -E allomorphs (Fig. 4A-B). Application of an anti-β_2_m mAb confirmed the particular recognition of assembled HLA-C and -G (Suppl. Fig. 3C-D). As seen before, the anti-HA antibody co-immunoprecipitated US10 with all tested HLA-I (Fig. 4A, C).

**Fig. 4.**
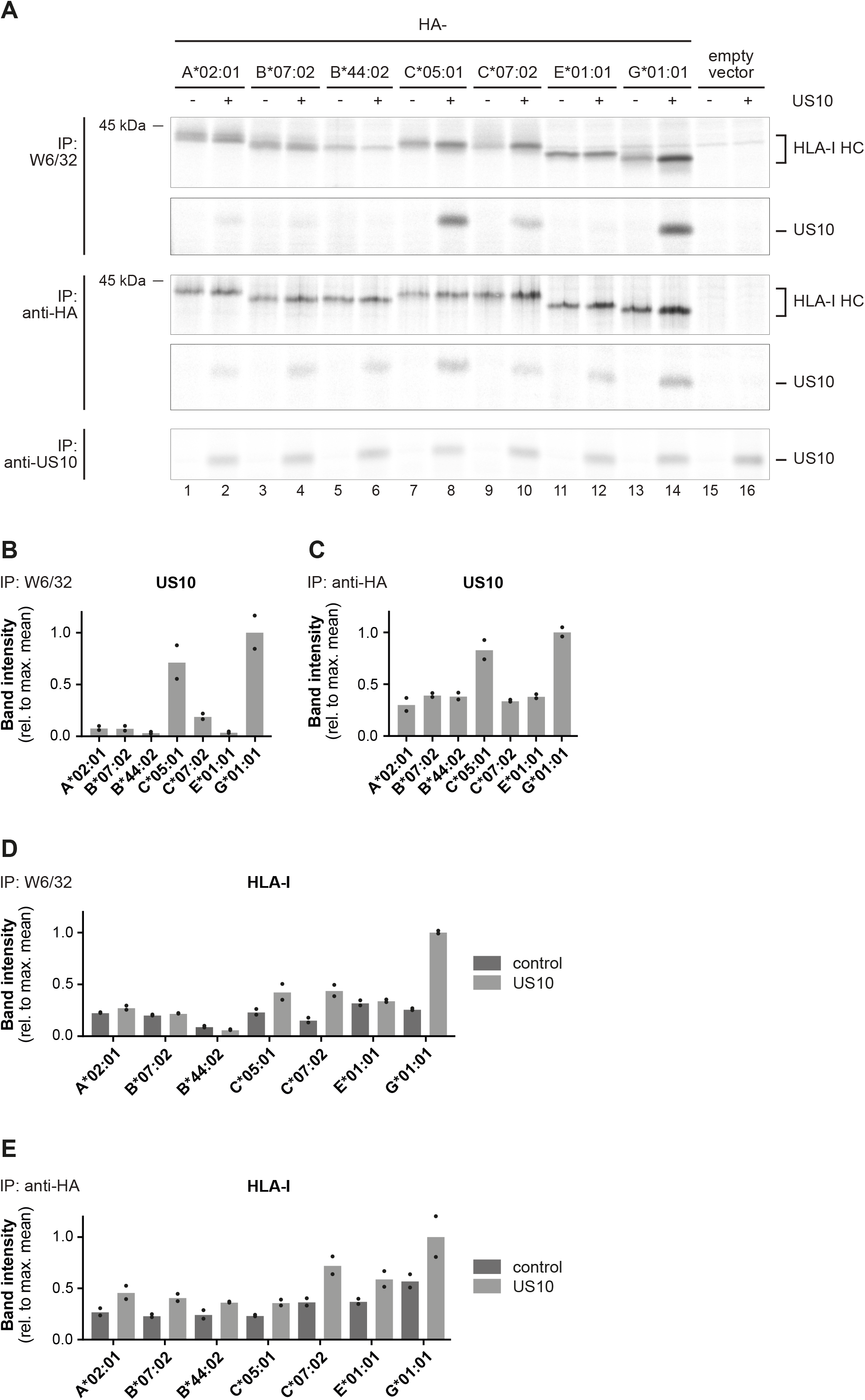
US10 selectively interacts with HLA-C and -G in their assembled forms. (A) HLA-I knock-out HeLa cells were transiently co-transfected with indicated HA-HLA-I-expressing plasmids comprising a mutated gRNA binding site together with a US10- or a control-pIRES-EGFP plasmid. To improve assembly of HLA-E, UL40 (comprising an HLA-E ligand) was expressed with HLA-E. Cells were metabolically labeled for 2h and immunoprecipitations were performed as in Fig. 1C; antibodies were applied as indicated to the left. (B-C) Relative signal strengths from single bands of US10 in the W6/32 (B) and anti-HA (C) immunoprecipitation samples are shown. Dots represent individual values and bars mean values thereof from two independent experiments (biological replicates). (D-E) Relative signal strengths from single bands of HLA-I HCs in the W6/32 (D) and anti-HA (E) immunoprecipitation samples are shown. Dots represent individual values and bars mean values thereof from two independent experiments (biological replicates).

Interestingly, in samples where US10 was strongly co-immunoprecipitated with the mAb W6/32 also induced HC/β_2_m assembly was observed, particularly pronounced for HLA-C*05:01, -C*07:02, and - G*01:01. In contrast, the tapasin-dependent HLA-B*44:02 allotype showed a reduced level of β_2_m assembly in the presence of US10 (Fig. 4D-E). The β_2_m-assembly with other HLA-I molecules was not affected by US10.

In conclusion, US10 possesses a particular specificity for HLA-C and -G. It promotes their assembly with β_2_m and forms a stable complex with HC/β_2_m heterodimers.

### Specialization of HCMV-encoded HLA-I immunoevasins

To gain insights into the individual roles of HCMV-encoded HLA-I immunoevasins, we next assessed HLA-I allotype specific effects exerted by single HCMV-encoded HLA-I immunoevasins. We compared the effects of US2, US3, and US11 on a selected panel of HLA-I molecules, as described for US10 in Fig. 1A. The immunoevasins displayed distinct profiles of susceptible and resistant HLA-I molecules (Fig. 5A). In accordance with our recent study (Zimmermann et al., 2019), US11 strongly reduced the expression of HLA-A*02:01 but not of HLA-B allotypes (Fig. 5A-B). US2 also effectively targeted HLA-A*02:01, and, somewhat less, HLA-B molecules with HLA-B*07:02 being the most resistant HLA-B allotype (Fig. 5A-B), in concordance with previous observations (Barel et al., 2003b). Notably, for some HLA-I molecules, co-transfection of US3 resulted in two distinct cell populations regarding surface HLA-I levels. A more pronounced downregulation of HLA-I was observed in the EGFP^dim^ than in the EGFP^bright^ population (Fig. 5C and Suppl. Fig. 4A-B). We speculate that alternative splicing, the activity of which can be dependent on the strength of transcription (Saldi et al., 2016), gives rise to one or more US3 versions, leading to various effects on HLA-I. Yet, some HLA-I allotypes were strongly downregulated by US3 independently of the strength of the EGFP expression: HLA-B*44:02, -B*08:01, -C*04:01, -C*12:03, and -G*01:01. Interestingly, these are highly tapasin-dependent molecules. Indeed, the sensitivity towards US3 (EGFP^bright^ gate) correlated strongly with HLA-I tapasin dependency (Suppl. Fig. 4C), which is in agreement with the finding that US3 blocks tapasin function (Park et al., 2004). Therefore, regarding HLA-B, the inhibitory function of US3 congruently overlapped with that of US10, while both immunoevasins showed striking differences regarding the inhibition of HLA-C and HLA-E (Fig. 5D-E).

**Fig. 5.**
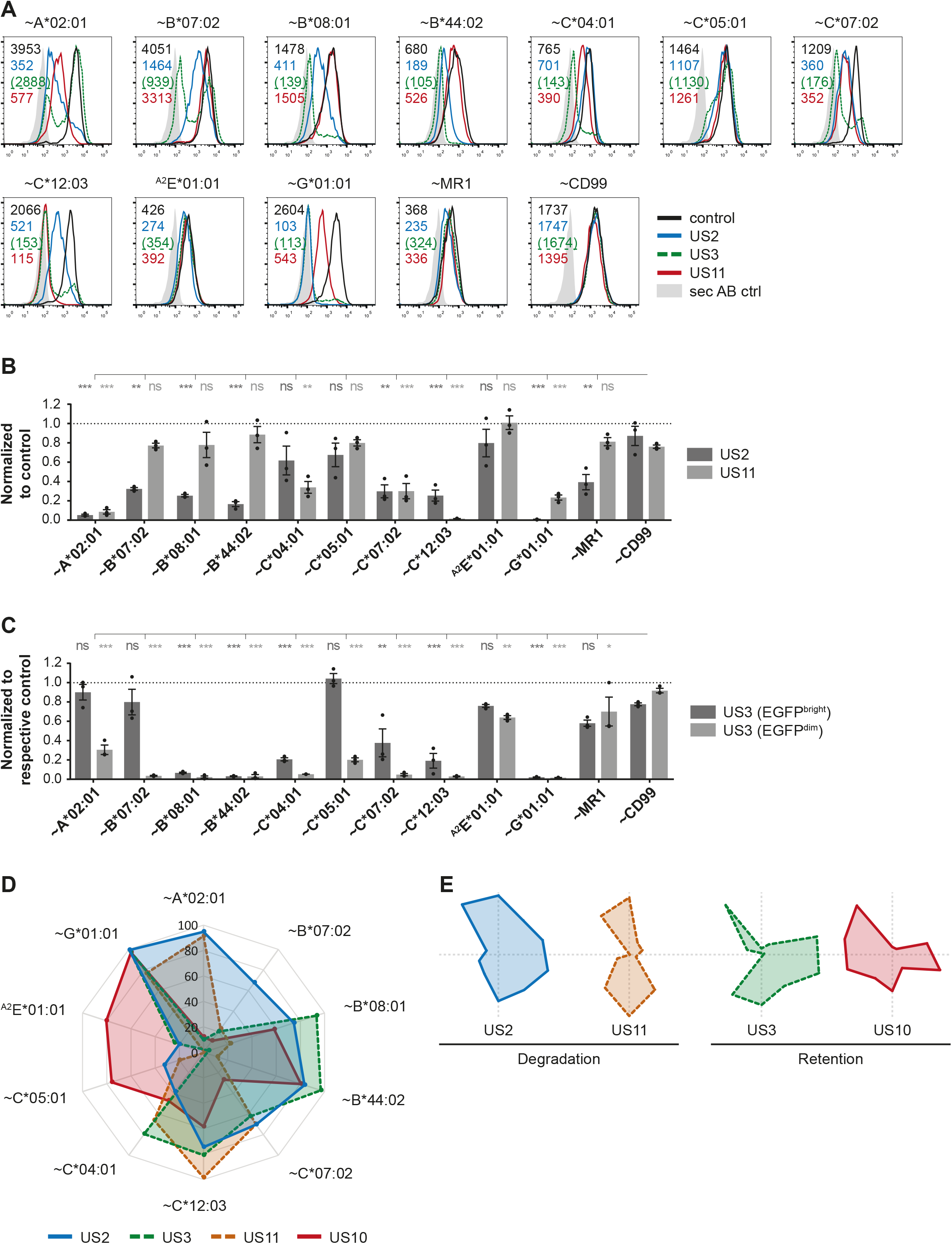
HLA-I allomorph dependent regulation by HCMV-encoded HLA-I immunoevasins. (A) HeLa cells were transiently co-transfected with HA-tagged molecules or untagged HLA-^A2^E*01:01 together with US2, US3, US11 or a control in the pIRES-EGFP plasmid. Cell surface expression was measured by flow cytometry using anti-HA or -HLA-E mAb (EGFP positive gate, gating as shown for US2 in Suppl. Fig. 4A). Representative histograms are shown. (B-C) Fold change by US2 and US11 (B) and US3 (C) was calculated by normalization of the MFI from (A) compared to control cells. Dots represent individual values and bars mean values ± SEM from three independent experiments (biological replicates). Significance compared to the individual HA-CD99 control was calculated using one-way ANOVA followed by Dunnett’s multiple comparison test. Fold change by US3 was determined in both EGFP^bright^ and EGFP^dim^ cells; gating and histograms are shown in Suppl. Fig. 4A-B. (D-E) Radar chart showing the susceptibility of HLA-I allomorphs to US2, US3, US11, and US10 (data from Fig. 1B and 5B-C shown as percentage downregulation). Susceptibility is given as percent loss of expression. Single profiles are shown in (E).

In our analysis, HLA-G*01:01 was highly susceptible to all immunoevasins, supporting previous findings for US2 and US3 (Barel et al., 2003b; Jun et al., 2000), while contrasting a study using cells stably expressing US11 (Barel et al., 2003a). This discrepancy could be due to cell type and expression differences. Regulation of the group of HLA-C allotypes varied strongly among the immunoevasins, possibly. Interestingly, HLA-C*05:01 which was both downregulated and strongly bound by US10, escaped all other immunoevasins. The same was true for HLA-E, which was effectively downregulated by US10, but not by the other immunoevasins, as has been shown before (Llano et al., 2003). These findings suggest that these HLA-I allotypes compose a specific niche for selective US10 modulation.

### Complex transcriptional regulation of US10 and US11

Next, we sought to validate the results regarding US10 modulation of HLA-I in HCMV-infected cells. We wanted to apply US10 siRNA to block US10 expression. However, previous Northern blot analysis reported two mRNAs transcribed from the *US11*/*US10* genome unit (Jones and Muzithras, 1991): A long mRNA starts upstream of the *US11* ORF encoding both US10 and US11, and a short mRNA starting in between *US11* and *US10* and comprising the *US10* ORF only. To reassess this using high-throughput sequencing techniques, we extracted *US10* and *US11* data from a recent meta-analysis of HCMV gene expression (Jürges et al., 2022). In this analysis, two transcription start site (TSS) profiling approaches, dRNA-seq (Sharma and Vogel, 2014; Whisnant et al., 2020) and STRIPE-seq (Policastro et al., 2020), were combined with metabolic RNA labeling (Erhard et al., 2019; Jurges et al., 2018) at multiple time points post-infection (p.i.) with HCMV. With basepair resolution, transcription start sites were pinpointed and their temporal expression kinetics was analyzed in detail. These data were integrated with previous Ribo-seq (Stern-Ginossar et al., 2012) and proteomics data (Weekes et al., 2014) as well as analysis of sequence motifs. Within the *US11*/*US10* genome unit (Fig. 6A), we found a single polyadenylation (poly-A) signal directly downstream of the *US10* ORF and two active TSSs located directly upstream of the *US10* and the *US11* ORFs, both having an upstream canonical TATA box at the expected distance (*US10* TSS: 31bp, *US11* TSS: 28bp; Fig. 6A). These TSSs are therefore consistent with the short and long mRNAs found by Northern blots.

**Fig. 6.**
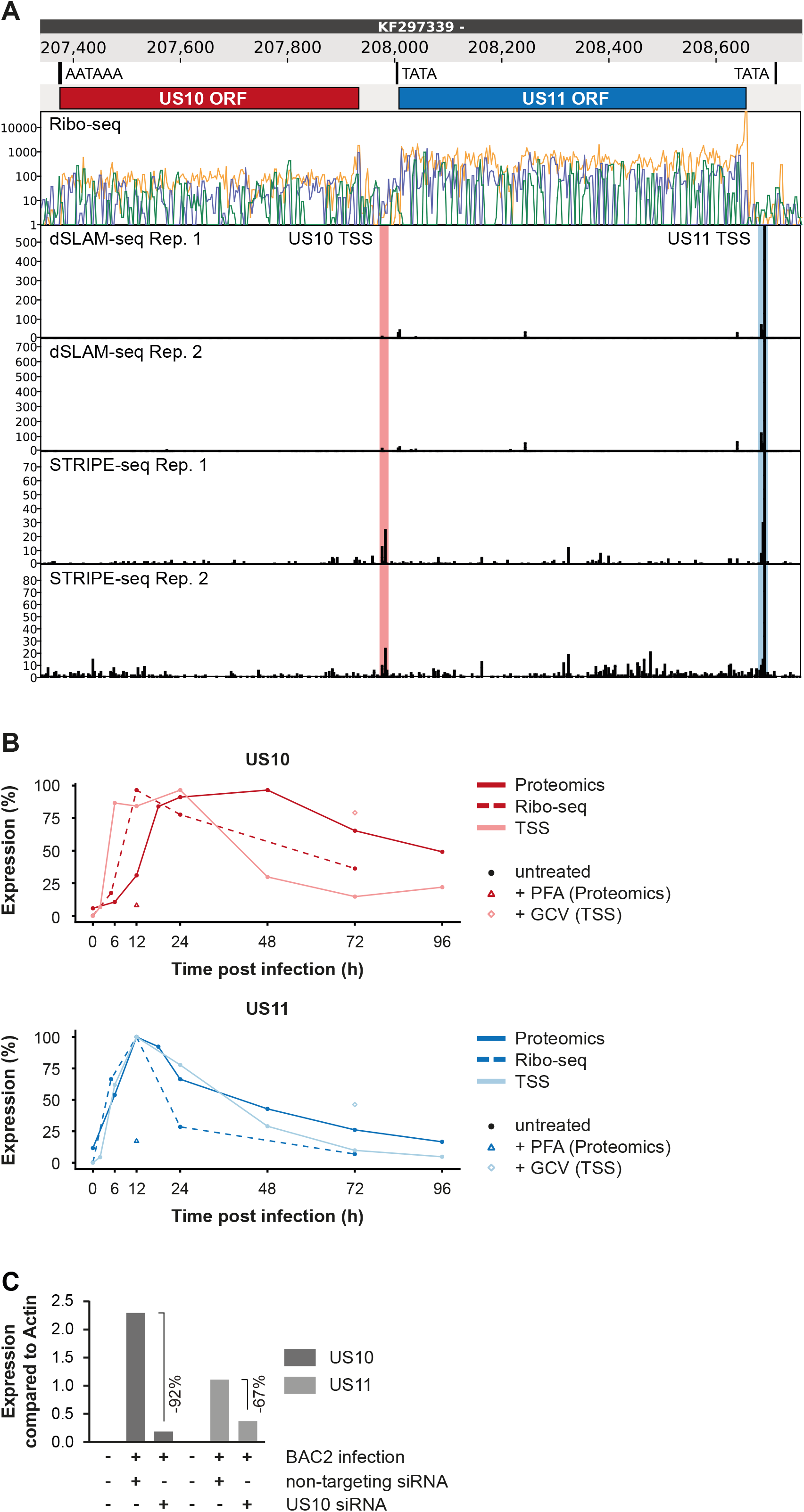
Analysis of *US10* and *US11* transcription and expression. (A) Genome browser showing the US10/11 locus. The tracks show (from top to bottom) the genomic position, identified sequence motifs, the known open reading frames (ORFs) encoding US10 and US11, the Ribo-seq signal mapped to the three possible reading frames (orange, green, blue) and the 5’ end read counts for TSS profiling data. The two identified TSS are indicated. (B) Time course plots depicting the temporal expression dynamics of US10 (top) and US11 (bottom). TSS profiling, Ribo-seq and proteomics data were scaled to the corresponding maximal value. Samples that were treated with genome replication inhibitors (PFA or GCV) are indicated. (C) MRC-5 fibroblasts were nucleofected with US10-specific or non-targeting siRNA 24 h prior to mock treatment or infection with HCMV Δ*US2-6* mutant BAC2 at an MOI of 5. At 24h p.i., RNA was isolated. Subsequently, cDNA was generated and analyzed by qPCR. Expression of US10 and US11 is shown compared to expression of actin.

The temporal kinetics of both TSSs were highly similar with steeply increasing expression at 6 h p.i. and slow decline after 24 h p.i. (Fig. 6B), but the activity of the *US11* TSS was much stronger (>20-fold) than the *US10* TSS. In agreement with this result, the Ribo-Seq data showed that translation of *US10* was at least one order of magnitude weaker than *US11*. Interestingly, a stronger US11 expression could be required to compensate for the short half-life of US11 (Baker and Tortorella, 2007), while lower expression level of US10 may suffice in combination with its slow turnover (Suppl. Fig. 5A). Taken together, these data provide evidence for a strongly expressed long mRNA expressing *US11* but also containing the *US10* sequence, and a weakly expressed short mRNA encoding *US10*. Furthermore, the distinct temporal kinetics of translation indicates that US11 and US10 are not polycistronically expressed from the long mRNA.

To test whether siRNA targeting of the *US10* sequence would also affect *US11*, we treated MRC-5 fibroblasts with US10 siRNA (Suppl. Fig. 5B-E) prior to HCMV infection and quantified *US10* and *US11* transcripts by qRT-PCR at 48h p.i. (Fig. 6C). The level of *US10* mRNA was reduced by 92% and US11 by 67%, verifying that *US11* is expressed from a transcript including the *US10* sequence.

### US10-specific siRNA has little effect on HLA-A, whereas HLA-B is strongly restored by the treatment

To measure the effect of US10 siRNA treatment on HCMV-infected MRC-5 fibroblasts, HLA-I surface expression was monitored. At 48h p.i., HLA-A*02:01 surface expression was not affected by this treatment, regardless whether the genes *US2-US6* were deleted (AD169VarL derived BAC mutant, BAC2 (Le-Trilling et al., 2020; Le et al., 2011)) or not (wild-type AD169VarL) (Fig. 7A and Suppl. Fig. 6A). This showed that despite a partial knock-down, US11 was able to control HLA-A*02:01 expression. In addition, this indicated that US10 had no significant effect on HLA-A*02:01 in HCMV-infected cells. Opposite to this, a clear upregulation of the HLA-B allotypes HLA-B*07:02 and - B*44:02 was observed in ΔUS2-US6-infected cells as a result of the US10 siRNA treatment. In previous experiments, HLA-B*07:02 was resistant to both US10 (Fig. 1) and US11 (Fig. 5), which could point either at an altered function in infection or at a co-regulation of HLA-B allotypes by US10 and US11. Not surprisingly, the presence of the genes *US2, US3*, and *US6* in HCMV wildtype infected cells blunted the effects of US10 siRNA on HLA-B allotypes. Importantly, the US10 siRNA effects on HLA-B*07:02 could be recapitulated by measuring activation of pp65/HLA-B*07:02-restricted CD8^+^ T-cells (Fig. 7B and Suppl. Fig. 6B), underlining the importance of the US11/US10 transcript expression for HLA-B antigen presentation.

**Fig. 7.**
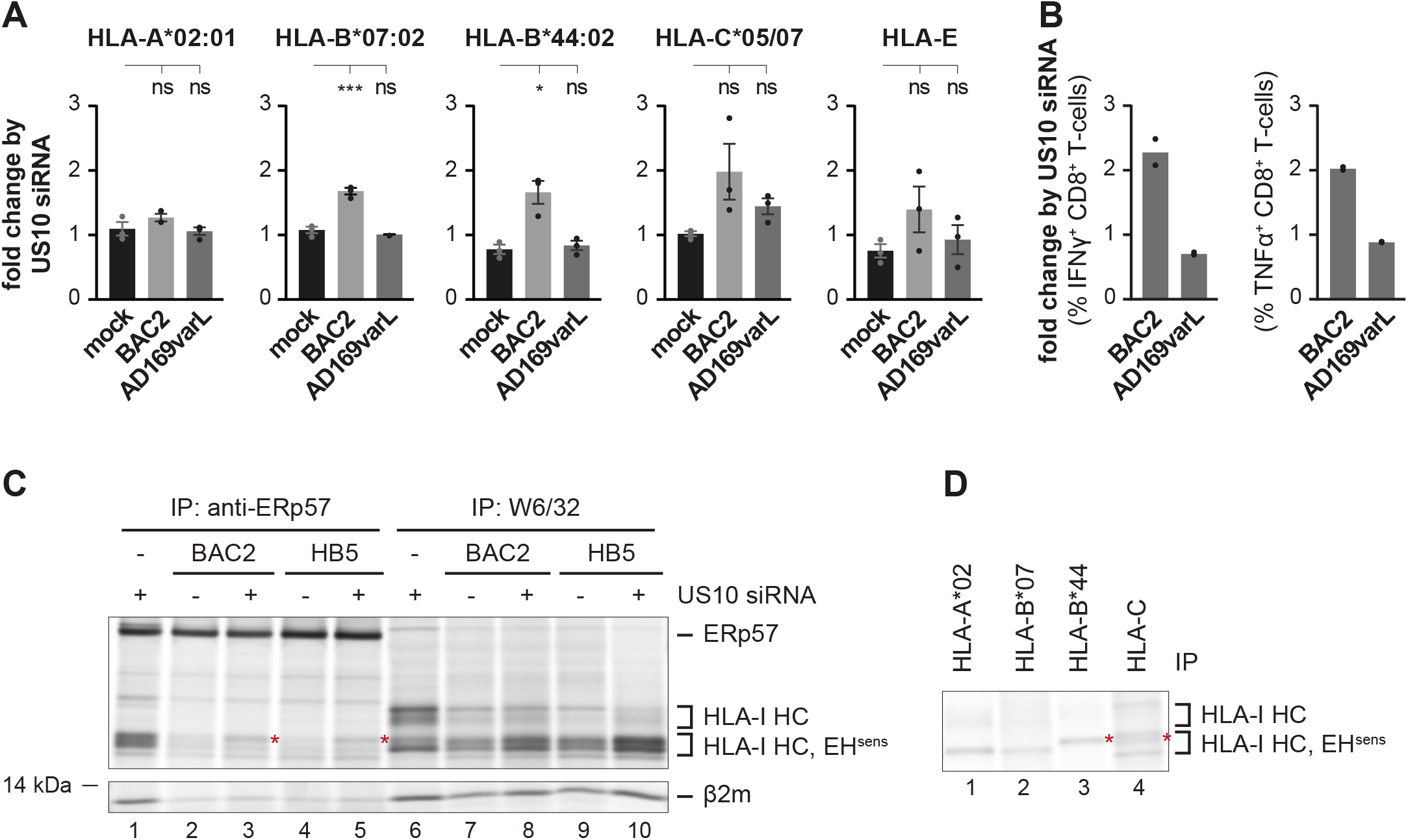
HLA-I regulation in US10 siRNA-treated HCMV-infected fibroblasts. (A) MRC-5 fibroblasts were nucleofected with US10-specific or non-targeting siRNA 24 h prior to mock treatment or infection at an MOI of 5 with HCMV Δ*US2-6* mutant BAC2 or with AD169varL. At 48h p.i., HLA-I surface expression was measured by flow cytometry using antibodies as indicated. Fold change by US10 was calculated as the ratio of the MFI of cells treated with US10 siRNA compared to NT treated cells. Dots represent individual values and bars mean values ± SEM from three independent experiments (biological replicates). Significance was calculated using one-way paired ANOVA followed by Dunnett’s multiple comparison test. (B) HFFα fibroblasts were treated and infected as in (A). At 24h p.i., the fibroblasts were co-cultured for five hours with HLA-B*07/pp65-restricted polyclonal CD8^+^ T-cells gained from PBMCs at an E/T ratio of 3:1. Activation of CD8^+^ T-cells was determined by intracellular IFNγ and TNFα stain. The percentage of IFNγ- or TNFα-expressing HLA-B*07/pp65 CD8^+^ T-cells was measured and the fold change by US10 siRNA was calculated. Dots represent individual values from two independent experiments (biological replicates)(representative dot plots are shown in Suppl. Fig. 6B). (C) MRC-5 fibroblasts were nucleofected with US10-specific or non-targeting siRNA 24h prior to mock treatment or infection with the HCMV Δ*US2-6* mutants BAC2 or HB5 at an MOI of 7. Proteins were metabolically labeled at 24h post-infection for 2h and immunoprecipitations using anti-ERp57 or W6/32 were performed. All samples were treated by EndoH. Asterisk: strongly increased HLA-I HC when applying US10 siRNA. One of two independent experiments (biological replicates) is shown. (D) BAC2 infected MRC-5 fibroblasts were treated as in (C) and HLA-I specific immunoprecipitations were performed as indicated. All samples were EndoH-treated prior to separation by SDS-PAGE.

To monitor HLA-C expression, we applied the mAb DT-9, which also recognizes HLA-E and therefore detects HLA-C*05:01, -C*07:02, and -E on MRC-5 fibroblasts. However, HLA-E surface expression determined by the mAb 3D12 remained low on MRC-5 cells also after HCMV infection (Suppl. Fig. 6A) and should only have a minor effect on the measured DT-9 signal. US10 siRNA treatment induced binding of DT-9 on both ΔUS2-US6- and wild-type-infected cells, but the effect was not significant (Fig. 7A).

In parallel to aforementioned flow cytometry analysis, we performed immunoprecipitations after metabolic labeling of MRC-5 fibroblasts infected with two different ΔUS2-US6 BAC mutants (BAC2 and HB5 [an AD169VarS-derived BAC mutant]). Treatment with US10 siRNA resulted in increased HLA-I assembly (Fig. 7C, lines 7-10), which most likely can be attributed to a reduction of US11 expression. Applying ERp57-antibodies, we analyzed the composition of the PLC and observed a clearly increased co-immunoprecipitation of a slowly migrating HLA-I HC (Fig. 7C, lines 3 and 5, red asterisk), but not of faster migrating HCs. Since HLA-A*02:01 and HLA-B*07:02 migrate faster in SDS-PAGE than HLA-B*44:02 and HLA-C allotypes (Fig. 7D), this finding supports the notion of a less prominent effect of the US10 siRNA on recruitment of HLA-A*02:01 and HLA-B*07:02 (lower MHC-I HC in the IP) to the PLC than of HLA-B*44:02 or -C (upper MHC-I HC in the IP). This finding fits well with the observation that HLA-B*15:03/-C*12:03 HCs in particular were excluded from the PLC in US10-expressing HeLa cells (Fig. 2) and suggests that also in HCMV-infected cells US10 blocks interaction of specific HLA-I with the PLC.

Taken together, US10 siRNA treatment underlines the lack of HLA-A*02:01 regulation by US10 also in HCMV-infected cells. Moreover, it shows that expression of HLA-B allotypes is sensitive to changes in expression levels of US10 and/or US11.

## Discussion

In HCMV infection, HLA-I antigen presentation is targeted by a well-orchestrated temporal expression of multiple immunoevasins. This strategy based on distinct mechanisms allows HCMV to control HLA-I gene- and allotype-specific functions in a versatile way across multiple cell types and environmental conditions (Ahn et al., 1997; Halenius et al., 2015; Hengel et al., 1997; Jones and Sun, 1997; Jones et al., 1996; Lehner et al., 1997; Park et al., 2004; Prod’homme et al., 2012; Wiertz et al., 1996; Zimmermann et al., 2019). We have recently shown an explicit targeting of HLA-A locus products by US11, while HLA-B escapes US11-mediated degradation. The refined analysis suggests that even though HCMV-encoded immunoevasins are partly redundant in their functions, they also exhibit distinguished non-redundant preferences towards certain HLA-I geno- or allotypes. Here we show that US10 evolved to target a specific group of HLA-I molecules.

### HLA-I downregulation by US10 is HLA-I gene locus- and allotype-specific

To analyze the effect of US10 on HLA-I, we selected a panel of classical and non-classical HLA-I molecules, including killer cell immunoglobulin-like receptor (KIR) ligands (possessing Bw4, C1, and C2 epitopes) and molecules that are not recognized by KIRs. While a broad spectrum of regulation by US10 was revealed, a conclusive pattern correlating with KIR binding was not observed, suggesting that US10 has not evolved to modify recognition of a specific type of KIR receptors. We observed efficient downregulation of some classical HLA-B and -C molecules and of the non-classical HLA-E and -G. In contrast, expression of HLA-A molecules was insusceptible to US10, also in HCMV-infected cells. This multitude of various US10 effects could explain why HLA-I cell surface regulation by US10 was not observed previously using the anti-pan-HLA-I mAb W6/32 (Ahn et al., 1997).

The absence of cell-surface regulation of HLA-A was not due to a lack of US10 recognition and binding, since the conformation-independent anti-HA antibody directed against tagged HLA-I demonstrated a remarkably conserved binding capacity of US10 to all tested HLA-I heavy chains (HLA-A, -B, -C, -E, and -G). However, consistent with unaffected cell surface expression of HA-tagged HLA-A allotypes, endogenous HLA-A*68:02 maturation as indicated by the gain of EndoH-resistant *N*-glycans, was only modestly delayed in HeLa cells. This suggesting that despite association with US10, HLA-A*68:02 can escape retention.

### Functional analysis of the PLC reveals HLA-I genotypic differences in US10 targeting

US10-mediated inhibition of HLA-B allotypes correlated strongly with their dependency on tapasin. Since HLA-A allotypes were not inhibited by US10, including the highly tapasin-dependent HLA-A*01:01, we consider it unlikely that US10 acts directly on tapasin. This notion was confirmed by additional downregulation of US10-sensitive HLA-I allotypes in tapasin-deficient cells (see Fig. 3C). While HLA-I interaction with the PLC was blocked in US10-expressing cells, this effect was less prominent for HLA-A*68:02. Our data support a model in which the affinity of US10 for HLA-A allotypes is lower than for HLA-B. This could explain a sufficient escape of HLA-A allotypes from US10 prior to entry in the PLC, or the acquisition of PLC-independent peptides which could be supported by TAPBPR preferentially associating with HLA-A (Boyle et al., 2013; Ilca et al., 2019).

In contrast to US2 (Suppl. Fig. 1), US3 (Park et al., 2004), and US11 (Zimmermann et al., 2019), US10 was not co-immunoprecipitated with the PLC. This suggests that US10 attacks HLA-I molecules in a way that precludes PLC integration. Hence, an efficient sequestration of tapasin-dependent HLA-B alloforms by US10 before entry in the PLC could largely compromise peptide-loading of certain HLA-B alleles such as HLA-B*44:02, while tapasin-independent -B*44:05 and -B*07:02 molecules might escape from US10 attack through acquiring tapasin-independent peptides followed by conformational maturation rendering such HLA-B allotypes US10-insensitive. Future studies will reveal whether US10 interference with PLC function will result in an aberrant HLA-I ligandome.

We have described a similarly impaired interaction of HLA-I with the PLC in HCMV-infected cells but did not assign this phenotype to a specific HCMV gene product (Halenius et al., 2011). Here, we showed that US10 contributes to this phenotype in HCMV-infected cells.

The unexpected finding that US10 associated with HLA-C and -G in their peptide-loaded, β_2_m-dimerized form much more efficiently than with HLA-A, -B, and -E suggests that US10 has evolved an independent molecular targeting strategy for HLA-C and -G. This presumption could explain why downregulation of HLA-C allotypes did not correlate with their level of tapasin dependency. Moreover, we noticed that US10 was able to induce β_2_m-dimerization of HLA-C and -G molecules. Admittedly, this stands in contrast to previous findings, demonstrating US10-facilitated degradation of HLA-G (Park et al., 2010). We have noticed that the US10 protein is highly sensitive to modifications (e.g. epitope tags) in regard to stability and localization and believe that changes in the US10 sequence may lead to aberrant effects on HLA-I. Therefore, in this study, we avoided the use of epitope tagged versions of US10.

Further, in contrast to HLA-C and -G, US10 reduced β_2_m-dimerization of the tapasin-dependent HLA-B*44:02, but not of the tapasin-independent HLA-B*07:02 allotype. These results indicate that binding of US10 and β_2_m to HLA-I heavy chains is not mutually exclusive and tapasin-independent peptide loading of HLA-A and particular HLA-B allotypes might represent a mechanism to escape US10 attack. Altogether, this illustrates the remarkable subtlety exerted by US10 interfering with the maturation of HLA-I molecules in both an HLA locus- and allotype-dependent manner. Hence, a gain in understanding of US10 selectivity may broaden our understanding of differences between HLA-I genotypes.

### HCMV-encoded HLA-I immunoevasins have both redundant and non-redundant specificities for HLA-I

In our comparative analysis of US2, US3, and US11, distinct profiles of HLA-I susceptibility for the individual immunoevasins were observed. These essentially covered all tested HLA-I molecules and only partially overlapped with US10. Notably, HLA-C*05:01 and HLA-E, the surface expression of which was strongly reduced by US10, were the most resistant to US2, US3, and US11 interference. While HLA-E is controlled by UL40 and US6 in HCMV-infected cells (Llano et al., 2003; Prod’homme et al., 2012; Ulbrecht et al., 2003), the importance of US10 in this context requires further investigation.

In accordance with previous findings (Park et al., 2004), we measured a highly significant correlation between HLA-I tapasin dependency and strength of US3-mediated downregulation, irrespective of the HLA-I genotype. This difference between US10 and US3 further highlights the distinct mechanisms of action of these two immunoevasins. Interestingly, some HLA-I allotypes, including HLA-B*07:02, showed increased resistance towards US3 in EGFP^bright^ HeLa cells (see Suppl. Fig. 4B). The *US3* gene encodes a splice variant lacking the transmembrane segment and this negatively affects US3 function (Jones et al., 1996; Shin et al., 2006). The *US3* cDNA used in this study comprises some of these splice sites and could produce a similar US3 variant without the transmembrane segment.

It is of particular interest that we observed highly variable allotype-dependent effects on HLA-C. Indeed, this appears to be a common feature of US2, US3, US10, and US11 and likely is connected to the explicit dual role of HLA-C allotypes in being ligands for both KIRs and αβ T-cell receptors (Djaoud and Parham, 2020). Long periods of co-evolution of virus and host may have led to a subtle equilibrium maintaining cell surface expression of a subset of HLA-C alleles. Despite frequent HLA-C*07:02-restricted HCMV/IE1-specific CD8^+^ T-cell responses (Schlott et al., 2018), HLA-C*07:02 expression could be beneficial for HCMV-infected individuals: it was shown that HLA-C*07:02 protects HCMV-infected cells from NK cell lysis in a KIR dependent manner (Ameres et al., 2013). A more comprehensive analysis of individual HLA-C allotypes regarding US10 susceptibility and comparison with HCMV infection rates and HCMV-specific CD8^+^ T-cell responses are required to gain more insight into this interesting topic. Moreover, a clear dissection of US10 and US11 effects in HCMV-infected cells was not possible due to loss of both mRNAs with the US10 siRNA. The generation of a set of HCMV mutants granting deletion of the selected gene while not changing transcription of the other, will be required to gain deeper understanding into how these genes are coordinated in infection to control HLA-I antigen presentation.

### A summary and model of US10-mediated HLA-I locus- and allotype-specific effects

US10 is able to bind to all HLA-I molecules early after their synthesis and prior to dimerization with β_2_m (Fig. 8). HLA-A overcomes this interaction in a manner that is not yet defined, but that might involve a timely displacement of US10 by binding to peptide and β_2_m. A lower affinity of US10 for HLA-A could be the major reason for this, as well as access to an alternative route for peptide loading, e.g. peptide loading facilitated by TAPBPR, a tapasin like chaperone (Thomas and Tampe, 2017).

**Fig. 8.**
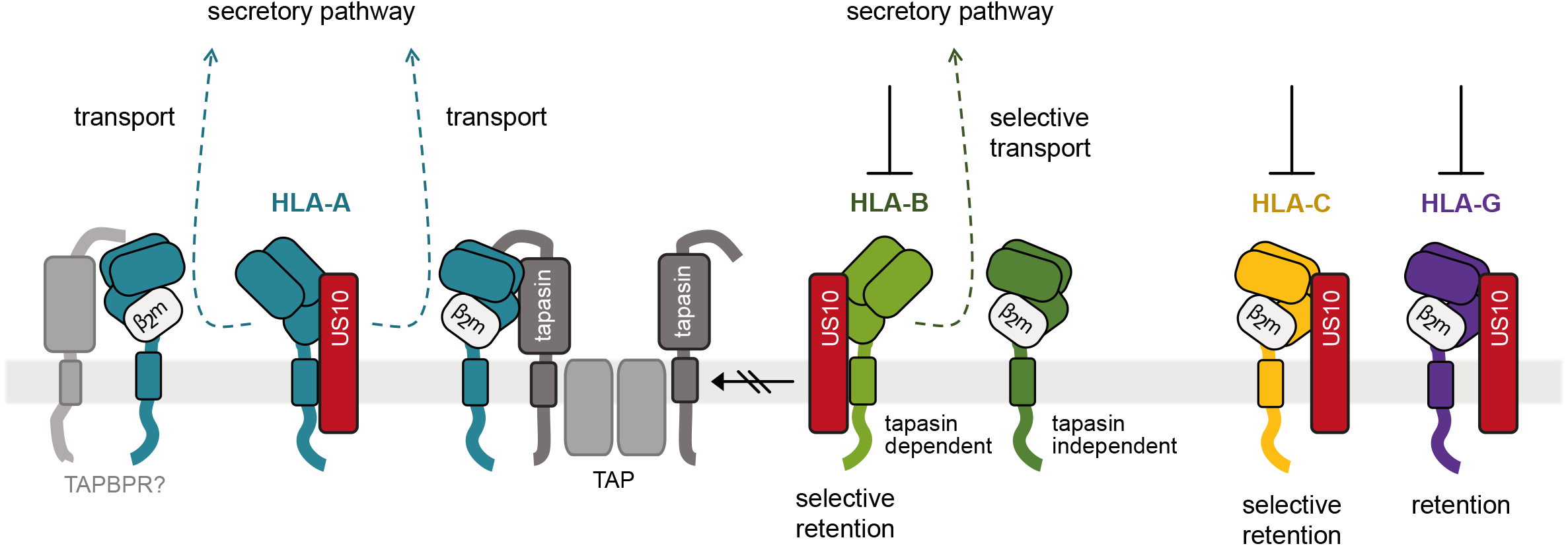
Model of HLA-I geno- and allotype-dependent targeting by US10. US10 is able to bind to all HLA-I HCs early after their synthesis. HLA-A (blue) overcomes this interaction by gaining access to tapasin or an alternative route for peptide loading, possibly TAPBPR. US10 binding to HLA-I prohibits recruitment to the PLC and tapasin-dependent HLA-B allotypes (light green) are therefore retained in the ER. Some HLA-B allotypes (dark green) can be loaded with peptides independently of tapasin enabling them to fold faster and escape US10. HLA-C (yellow) and HLA-G (violet) strongly associate with US10 also in a β_2_m-assembled conformation and are retained in the ER.

US10 binding to HLA-I prohibits recruitment to the PLC, which strongly affects cell surface expression of tapasin-dependent HLA-B allotypes. Possibly, the ability to be loaded with peptides independently of tapasin enables HLA-B molecules to fold faster and gain an assembled conformation that escapes US10. While the result of this is similar to HLA-A escape from US10, the molecular basis could be different due to the preference of TAPBPR for HLA-A allotypes (Ilca et al., 2019). An appealing idea is that, in HCMV infection, US10 may sort HLA-I prior to attack by US2, US3, or US11 to strengthen or dampen the effects, depending on the gene- or allotype.

Since opposite to HLA-A/-B/-E, HLA-C and -G can strongly associate with US10 also in a W6/32-reactive conformation, β_2_m- and peptide-binding apparently does not in general induce US10- resistant conformations. We predict a distinct strategy for US10 in manipulation of HLA-C and -G, which might depend on a small number of polymorphic amino acids. Given the relevance of congenital HCMV infections and the fact that trophoblasts do not express HLA-A and -B, but HLA-C, - G, and -E (Hackmon et al., 2017) the question arises whether US10 has evolved a strategy to manipulate HLA-I in a distinct manner at the interface between mother and fetus.

## Material & methods

### Molecular cloning

MHC-I and MHC-I like molecules were cloned into a Tpn-SP-pIRES-EGFP vector via PstI or NsiI and BamHI. This vector encodes the tapasin signal peptide and an HA-tag N-terminally to the insertion site for the MHC-I sequence. The HA-tagged constructs where then subcloned into the puc2CL6IP vector using NheI and BamHI (all primers in Table 1), which was used for expression. Sequences for HLA-I molecules were obtained from different sources: HLA-C*04:01 and -G*01:01 from cDNA prepared from JEG-3 cells (ATCC HTB-36); HLA-A*24:02, -C*05:01 and H-2K^b^ were ordered as gBlock gene fragments (Integrated DNA Technologies); HA-MR1 was purchased in the pcDNA3.1 vector (BioCat); HA-tagged MICA*004 was obtained from a previously published construct (Seidel et al., 2015). The sequence for HLA-E*01:01 was obtained as a gBlock gene fragment. To add the signal peptide of HLA-A*02:01, it was included into a primer. RL8, US2, US2HA, US10, US10HA, UL40 and US9HA were amplified from AD169 HCMV cDNA (or a previously described construct on the case of US3HA) and inserted into pIRES-EGFP or puc2CL6IP via NheI/BamHI and XhoI/BamHI (for US2HA), respectively. Not mentioned constructs have been previously described (Beutler et al., 2013; Zimmermann et al., 2019). For mutation of HLA-I gRNA target sites, the Q5 site-directed mutagenesis kit (New England BioLabs) was applied. Plasmids were amplified in *E. coli*, NEB5-alpha from New England Biolabs (C2987).

**Table 1:**
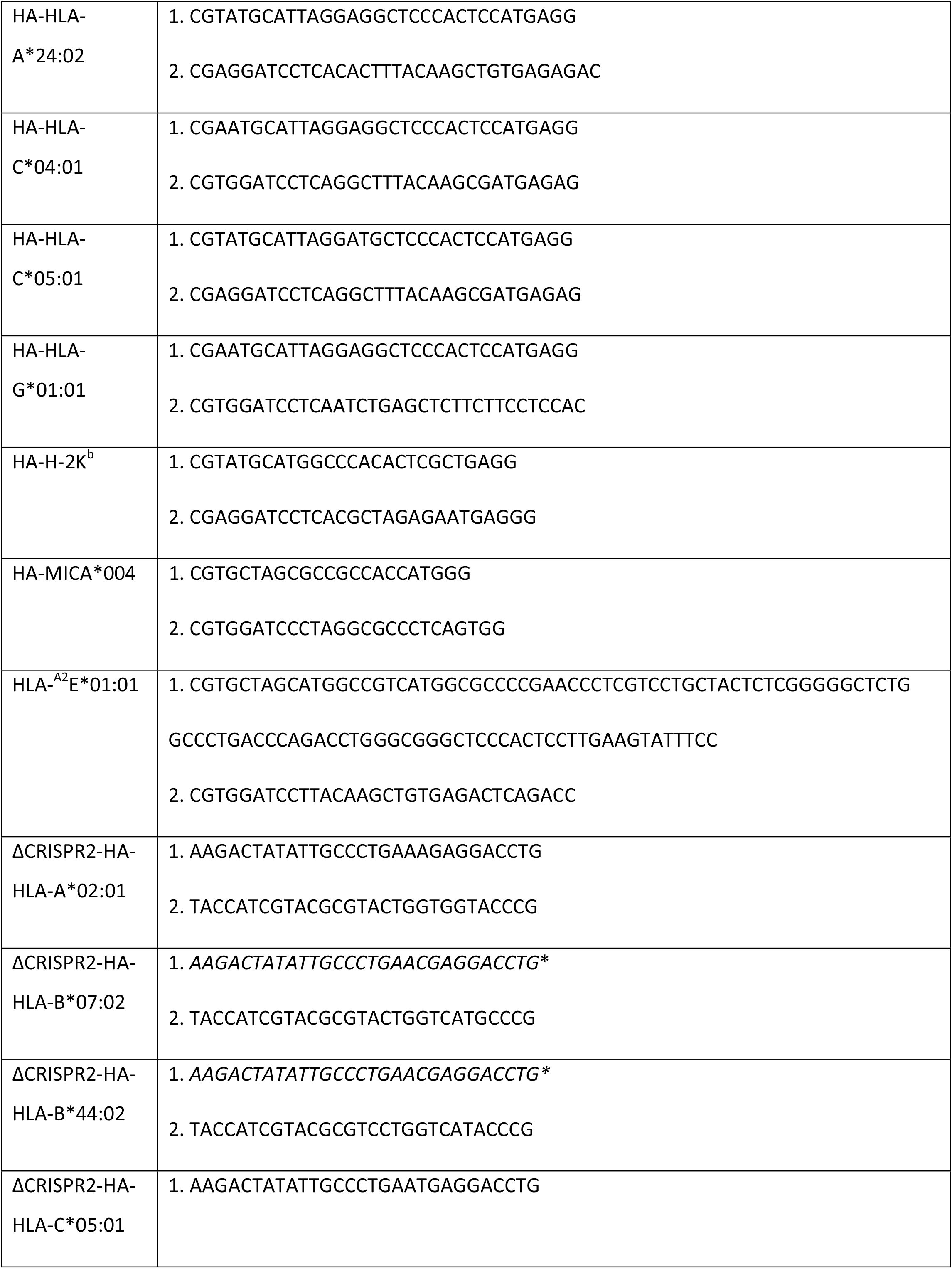
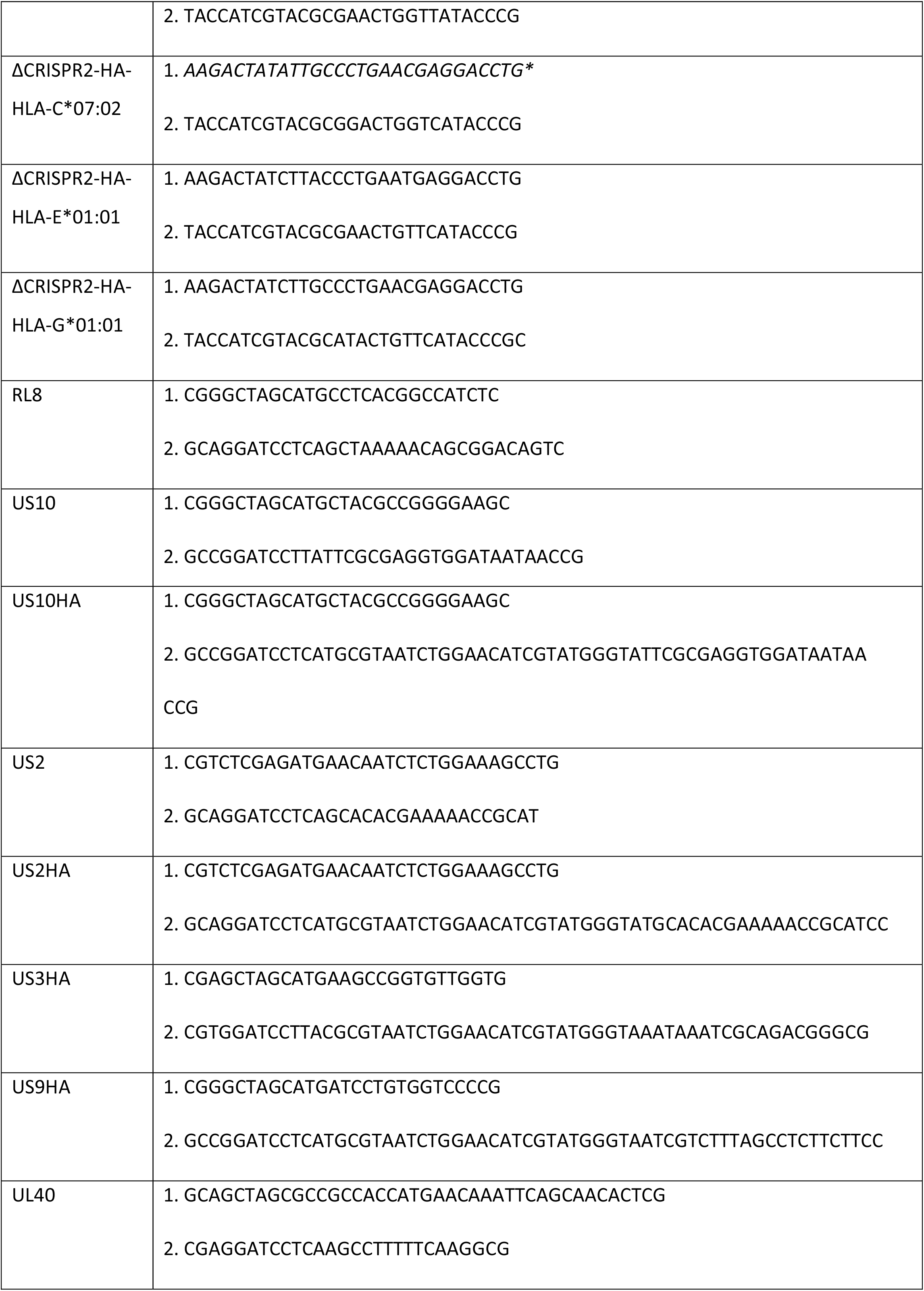
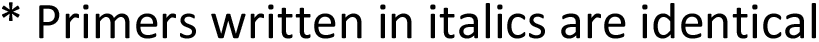
Primer sequences for molecular cloning.

### Cell culture, transfection, nucleofection and generation of stable cell lines

HeLa cells (human, ATCC CCL-2), MRC-5 fibroblasts (human, ATCC CCL-171), and HFFα human foreskin fibroblasts (a kind gift of Dieter Neumann-Haefelin and Valeria Kapper-Falcone, Institute of Virology, Medical Center University of Freiburg, Freiburg, Germany) were grown in DMEM (Life Technologies) supplemented with 10 % (v/v) FCS (Biochrom, Sigma-Aldrich or PAN-Biotech) and 1 % (v/v) penicillin/streptomycin (Life Technologies, stock: 5000 U/ml) at 37°C and 5 % CO_2_. HeLa cells stably expressing US9HA (control cells), US10 or US10HA were generated by lentiviral transduction. A tapasin knock-out cell line was generated by transient expression of SpCas9 and a corresponding gRNA targeting the sequence GCCCTATACGCCAGGCCTGG using plasmids kindly gifted by J. Keith Joung (Addgene plasmids #43861 and #43860) as detailed elsewhere (Muller et al., 2016). The HLA-I knock-out HeLa cells were generated as previously described (de Waard et al., 2021). For transient expression, HeLa cells were transfected with SuperFect (Qiagen) for 20h and fibroblasts cells were nucleofected using the SE Cell Line AD-Nucleofector Kit (Lonza). Knock-down procedures were performed with siRNA purchased from Riboxx (non-targeting siRNA: UUGUACUACACAAAAGUACCCCC; US10 targeting #1: UUCUGAAUAACACAGCCGCCCCC) applying the SE Cell Line Kit (Lonza) for fibroblasts and Lipofectamine RNAiMax (Invitrogen) for HeLa cells.

### Viruses

Generation, reconstitution, and propagation of the HCMV strain AD169VarL and the recombinant HCMV Δ*US2-6* mutants BAC2 (GenBank accession number MN900952.1) based on the AD169varL and HB5 based on the AD169VarS (GenBank accession number X17403) strains, were previously described (Borst et al., 1999; Halenius et al., 2011; Hengel et al., 1995; Le-Trilling et al., 2020; Le et al., 2011). In experimental settings fibroblasts were infected with an MOI of 5-7 with centrifugal enhancement (800 x g for 30 min).

### Antibodies

The following antibodies were used in this study: W6/32 (anti-pan-HLA-I assembled with β_2_m and peptide (Parham et al., 1979)), BB7.1 (anti-HLA-B*07 (Brodsky et al., 1979b)), BB7.2 (anti-HLA-A*02 (Brodsky et al., 1979b)), BBM.1 (anti-β_2_m (Brodsky et al., 1979a)), TT4-A20 (anti-B*44 (Tahara et al., 1990)), DT-9 (anti-HLA-C, BioLegend, 373302, RRID AB_2650941), 3D12 (anti-HLA-E, BioLegend, 342602, RRID AB_1659247), anti-HA produced in mouse (Sigma-Aldrich (H3663, RRID AB_262051)) or rabbit (Sigma-Aldrich (H6908, RRID AB_260070)), anti-ERp57 (abcam (ab13506, RRID AB_1140700), Millipore), anti-β-actin (Sigma-Aldrich, A2228, RRID AB_476697), anti-CD28 (BD Biosciences), APC-coupled anti-HA (Miltenyi Biotec (130-098-404, RRID AB_2751024)), APC-coupled IgG1-isotype control (Miltenyi Biotec, 130-113-200, RRID AB_2733881), APC-coupled anti-CD8 (BD Biosciences), APC-coupled anti-CD56 (Miltenyi Biotec (130-113-305, RRID AB_2726084)), APC-coupled goat anti-mouse IgG (BD Biosciences (550826, RRID AB_398465)), FITC-coupled anti-IFNγ (BD Biosciences, 554551, RRID AB_395473), PE/Cy7-coupled anti-TNFα (BioLegend, 506323, AB_2204356), HRP-coupled goat anti-mouse IgG (Dianova, 115-035-146, RRID AB_2307392), HRP-coupled goat anti-rabbit IgG (Sigma-Aldrich, 12-348, RRID AB_390191). Polyclonal anti-tapasin and anti-US10 were raised in rabbits (GenScript) using synthetic peptides (aa 418-428 and aa 54-67, respectively).

### Flow Cytometry

HeLa cells were harvested using trypsin and MRC-5 cells using accutase (Sigma-Aldrich). The cells were washed and stained with antibodies in PBS with 3 % (v/v) FCS. Cells were further stained with DAPI before flow cytometry analysis (FACS Canto II (BD Biosciences), FlowJo (Tree Star)). IFNγ-treatment was performed with 1000 U/ml (BioLegend) for 16 hours. FcR blockage (FcR blocking reagent, Miltenyi Biotec) and fixation in 4 % paraformaldehyde were performed for infected cells. For intracellular staining, Cytofix/Cytoperm from BD Biosciences was applied.

### Immunoprecipitation

Immunoprecipitations were performed as previously described (Halenius et al., 2011). Briefly, cells were cultured in 6-well plates and metabolically labeled (Easytag Express ^35^S-Met-Cys protein labeling mix (Perkin Elmer)) with 0.2 mCi/ml for various times. In pulse chase experiments, cells were washed and incubated in DMEM containing additional methionine/cysteine (Sigma-Aldrich) for different time periods. Cells were lysed in 1 % (w/v) digitonin (Calbiochem) lysis buffer (140 mM NaCl, 20 mM Tris (pH 7.6), 5 mM MgCl_2_) containing 1x cOmplete protease inhibitor (Roche). For immunoprecipitation with several antibodies, identical lysates were pooled before separation into aliquots for specific immunoprecipitations. Lysates were incubated with antibodies for one hour at 4°C in an overhead tumbler before immune complexes were retrieved by protein G (GE Healthcare or abcam) or A Sepharose (GE Healthcare). The beads were washed with increasing NaCl concentrations. Endoglycosidase H digestion was performed according to manufacturer’s instructions (New England BioLabs). Complexes were dissociated at 95°C in sample buffer containing 150 mM DTT prior to loading on a gradient SDS-PAGE. Gels were fixed, dried and exposed to phosphor screens analyzed by Typhoon FLA 7000 (GE Healthcare) or to x-ray films. For better illustration, contrast and light settings were adjusted in the figures. Band intensities were determined by the ImageQuant TL Software (GE Healthcare Life Sciences). The background signal was subtracted from values used for graphical visualization of quantification.

### Western blot analysis

Cell lysates were separated by SDS-PAGE and transferred to a nitrocellulose membrane (Amersham Protan, GE Healthcare). Incubation with specific antibodies was followed by peroxidase-conjugated secondary antibodies and detection using SignalFire ECL Reagent (Cell Signaling Technology) and the ChemiDoc XRS System (Bio-Rad).

### Transcriptional analysis of US10 and US11

Functional genomics data for the US10/11 locus were extracted from our recent integrative meta-analysis (Jürges et al., 2022). Briefly, primary human foreskin fibroblasts (HFFα) were infected with HCMV strain TB40 at MOI 10 and two transcription start site (TSS) profiling methods (dSLAM-seq (Sharma and Vogel, 2014; Whisnant et al., 2020), Stripe-seq (Policastro et al., 2020)) were applied. All TSS profiling data were analyzed using iTiSS (Jurges et al., 2021), published Ribo-seq data (Stern-Ginossar et al., 2012) was analyzed using PRICE (Erhard et al., 2018), and protein expression data were directly taken from the supplementary material of a previous study (Weekes et al., 2014). The genome browser to visualize all data sets is available on zenodo (https://doi.org/10.5281/zenodo.5801030) and a web-based tool for generating time course plots is available on the project web-site (https://erhard-lab.de/web-platforms).

### Quantitative RT-PCR

RNA was isolated using the NucleoSpin RNA kit from Macherey-Nagel according to manufacturer’s instructions. cDNA was generated using the QuantiTect Reverse Transcription Kit from Qiagen according to manufacturer’s instruction. The qPCR was performed with a SYBR Green PCR Master mix (Applied Biosystems) and the primers US10-SYBR1 ACGACGGGGAAAATCACGAA, US10-SYBR2 CAGAGTAGTTTCGGGGTCGG, US11-SYBR1 TTGTTCGAAGATCGCCGTCT, US11-SYBR2 AAAATGTCGGTGCAGCCAAC (Suppl. Fig. 5B) and Qiagen control primer Hs_ACTB_1_SG QuantiTect. Expression of US10 and US11 was determined compared to control using the 2^-ΔCt^ method.

### Generation of HLA-B*07/pp65-restricted CD8^+^ T-cell polyclones

PBMCs from an HCMV-seropositive, healthy, HLA-B*07:02-positive donor (female) were gained from EDTA blood by density separation (Lymphocyte Separation Medium (AnproTech)) and CD8^+^ T-cells were isolated with the human CD8^+^ T-cell isolation kit (Miltenyi Biotec) and a MidiMACS LS-column according to manufacturer’s instructions. Isolated CD8^+^ T-cells (2x 1.5×10^6^ cells) or original PBMCs (2x 2×10^6^ cells) were cultured in RPMI 1640 medium (supplemented with 10 % (v/v) FCS (PAN Biotech), 1 % (v/v) penicillin/streptomycin (Life Technologies) and 1.5 % (v/v) HEPES (Life Technologies)). 0.5 µg/ml anti-CD28 antibody (BD Biosciences) and 5 µM of an HLA-B*07:02-specific pp65 peptide (417-426): TPRVTGGGAM (Genaxxon (purity: 98.3 %)) were added to the medium. 50 % of the medium was replaced by fresh, supplemented RPMI with 20 IU/ml IL-2 (Stemcell) every three days. After 14 days, specificity of the cells was tested by staining with tetramers generated from HLA-I easYmer and streptavidin-PE (BD Biosciences) according to the manufacturer’s instructions (immunAware). For analysis, 5×10^4^ cells were incubated with the tetramer, further stained with anti-CD8 (BD Biosciences) and viability dye (EBioscience), washed, fixed in 2 % PFA and measured on a FACS Canto. The project has been approved by the ethical committee at the Albert-Ludwigs-University Freiburg (22-1196).

### Analysis of activation of HLA-B/CMV-restricted CD8^+^ T-cells by HCMV infection

1×10^5^ HFFα fibroblasts nucleofected with US10- or non-targeting siRNA were seeded in DMEM (Life Technologies) supplemented with 10 % (v/v) FCS (PAN Biotech) and 1 % (v/v) penicillin/streptomycin (Life Technologies). After 24 hours, cells were infected with HCMV at an MOI of 5. At 24h p.i., the infected cells were co-cultured with HLA-B/CMV-restricted CD8^+^ T-cells for five hours (E:T ratio: 3:1) in RPMI 1640 medium (Life Technologies)(supplemented with 10 % (v/v) FCS (PAN Biotech), 1 % (v/v) penicillin/streptomycin (Life Technologies) and 1.5 % (v/v) HEPES (Life Technologies)). GolgiStop and GolgiPlug (BD Biosciences) were used according to manufacturer’s instructions and added to the co-culture. Activation of the CD8^+^ T-cells was analyzed by flow cytometry. T-cells were transferred to a 96 well plate after co-culturing and surface staining of CD8 (BD Biosciences) was performed. Additionally, a viability dye-Cyan (EBioscience) was added. After fixation and permeabilization (BD Cytofix/Cytoperm Fixation/Permeabilization Kit), intracellular staining with FITC-coupled anti-IFNγ (BD Biosciences) and PE/Cy7-coupled anti-TNFα (BioLegend) was performed. Cells were measured on the FACS Canto (BD Biosciences). As a positive control, T-cells were cultured with 15 µM HLA-B-specific CMV peptide for five hours.

### Statistics

Calculations based on flow cytometric results were performed from the median fluorescence intensity (MFI) after subtraction of background signal (secondary antibody or isotype control) and shown with the standard error of the mean (SEM). For statistical analysis of the siRNA infection experiments and of the allotype-specific regulation of HLA-I surface expression after transient transfection, one-way ANOVA followed by Dunnett’s multiple comparison test was performed as recommended by the GraphPad Prism Software (v8). The correlation analysis (two-tailed) was performed with the same software. A p-value < 0.05 was considered significant (*, p < 0.05; **, p < 0.005; ***, p < 0.0005). The radar chart was generated using Microsoft Excel (2016).

### Data and code availability

Already published data used for our analysis can be found at these sites: dRNA-seq and STRIPE-seq data at NCBI Gene Expression Omnibus, GEO (accession number GSE191299), PRO(cap)-seq at GEO (GSE113394), and Ribo-seq at GEO (GSE41605), PacBio and MinION at the European Nucleotide Archive (accession number PRJEB25680), and protein expression data from the supplementary material of the reference (Weekes et al., 2014). These have been integrated into a genome browser and a web-based visualization platform (https://doi.org/10.5281/zenodo.5801030 and https://erhard-lab.de/web-platforms) PRICE is available at https://github.com/erhard-lab/price, the gedi toolkit at https://github.com/erhard-lab/gedi and iTiSS at https://github.com/erhard-lab/iTiSS. The MetagenePlot, a module for gedi, is available at https://github.com/erhard-lab/MetagenePlot. The source code of the gedi toolkit is available at https://github.com/erhard-lab/gedi. The source code for additional custom scripts can be found at Zenodo (https://doi.org/10.5281/zenodo.5801030).

### Materials availability

Requests for resources should be directed to the corresponding author.

## Acknowledgements

We are thankful for technical support by Magdalena Schwarzmüller. The work was supported by the Deutsche Forschungsgemeinschaft through the FOR2830 (HA 6035/2-2, ER 927/1-2, and DO 1275/7-2).

## Supplementary Figure Legends

**Suppl. Fig. 1.**
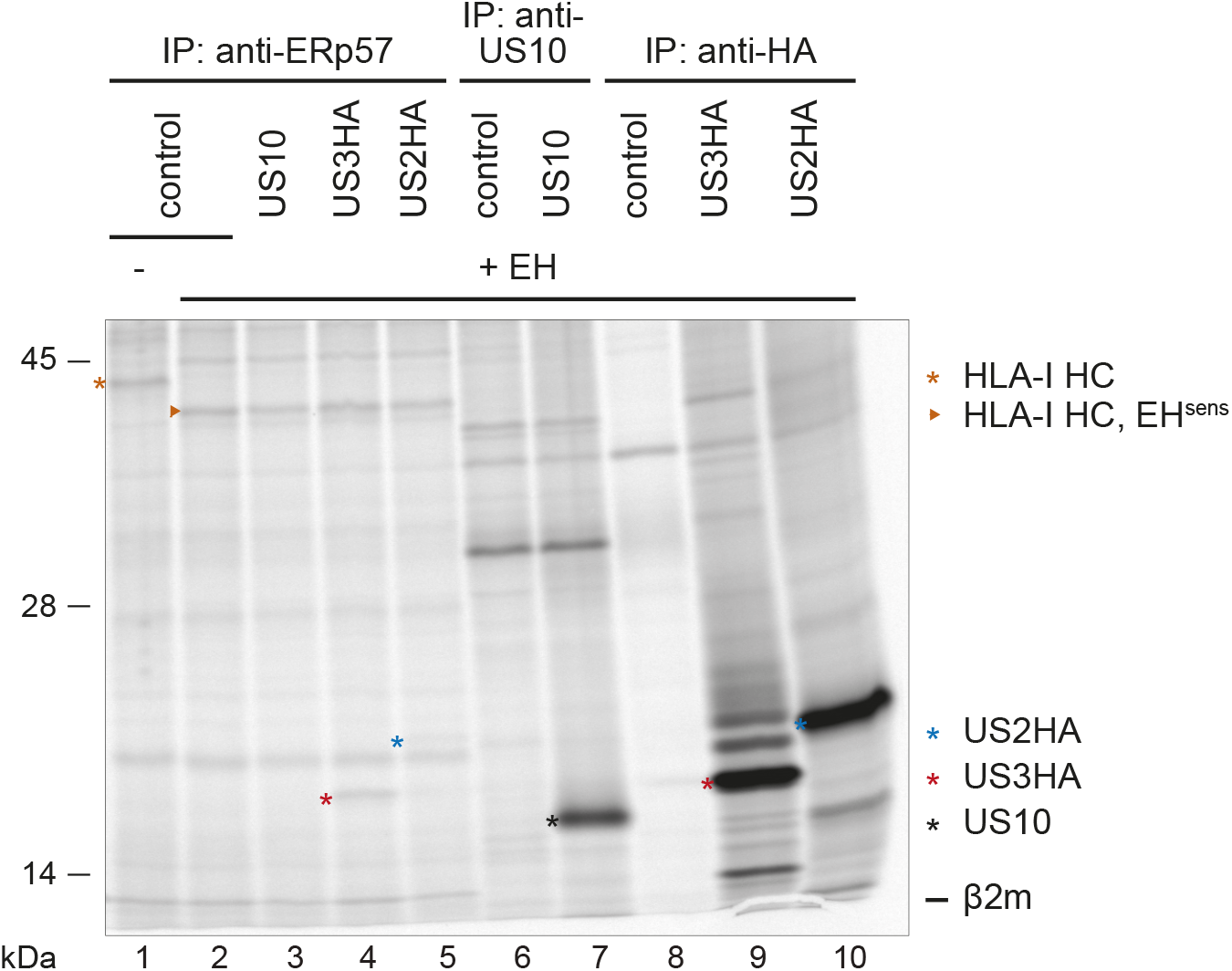
US10 is not co-immunoprecipitated with the PLC. (A) HeLa cells were transiently transfected with US10 (black asterisk), US3HA (red asterisk), US2HA (blue asterisk) or a control. At 20h post-transfection, proteins were metabolically labeled for 2h and immunoprecipitations using anti-ERp57, anti-HA or anti-US10 antibodies were performed and subjected to EndoH digest where indicated. Samples were separated by SDS-PAGE and labeled proteins were detected by autoradiography. One of two independent experiments (biological replicates) is shown.

**Suppl. Fig. 2.**
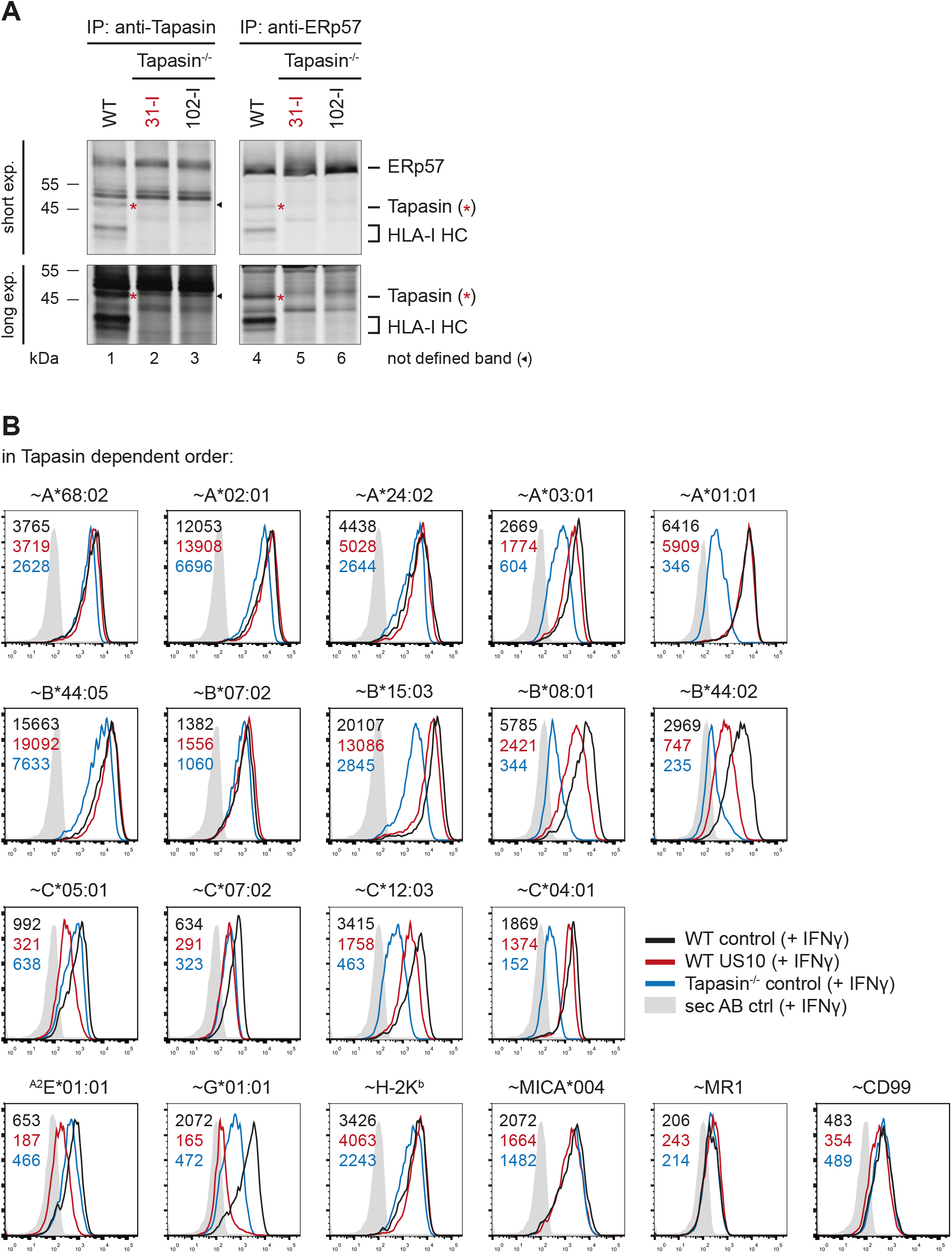
Measurement of tapasin independence and US10 resistance. (A) Wild-type HeLa cells and two clones of tapasin knock-out HeLa cells (31-I and 102-I) were metabolically labeled for 2h. Immunoprecipitations using anti-tapasin and anti-ERp57 were performed. Red asterisk: tapasin. Black arrow: unspecific band. A short and a long exposure are shown. Tapasin knock-out clone 31-I was used for all subsequent experiments. (B) Representative histograms for Fig. 3A.

**Suppl. Fig. 3.**
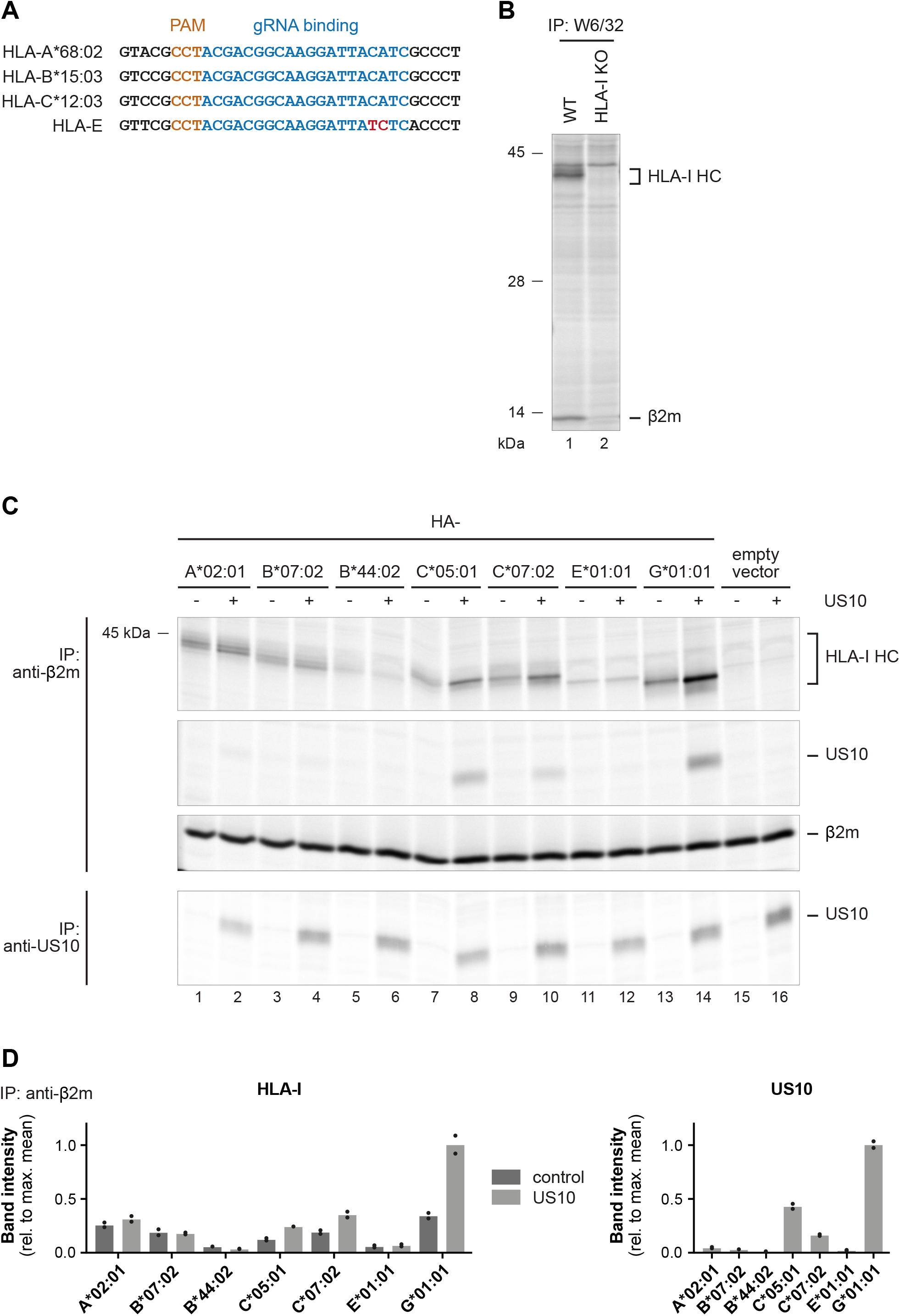
US10 selectively interacts with HLA-C and -G in their assembled conformation. (A) Alignment of the sequences of HeLa HLA-I (HLA-A*68:02, -B*15:03, -C*12:03 and -E). Highlighted are the PAM (orange) and the gRNA binding (blue) sites used for generation of the HLA-I knock-out HeLa cells. A mismatch in the HLA-E sequence is indicated in red. (B) Wild-type and HLA-I HeLa knock-out cells were metabolically labeled for 2h prior to an immunoprecipitation using W6/32. Retrieved proteins were separated by SDS-PAGE and detected by autoradiography. One of two independent experiments (biological replicates) is shown. (C) Immunoprecipitation was performed as described in Fig. 4A, but by applying the antibodies anti-β_2_m and anti-US10. (D) Relative signal strengths from single bands in the anti-β_2_m immunoprecipitation samples are shown. Dots represent individual values and bars mean values thereof from two independent experiments (biological replicates).

**Suppl. Fig. 4.**
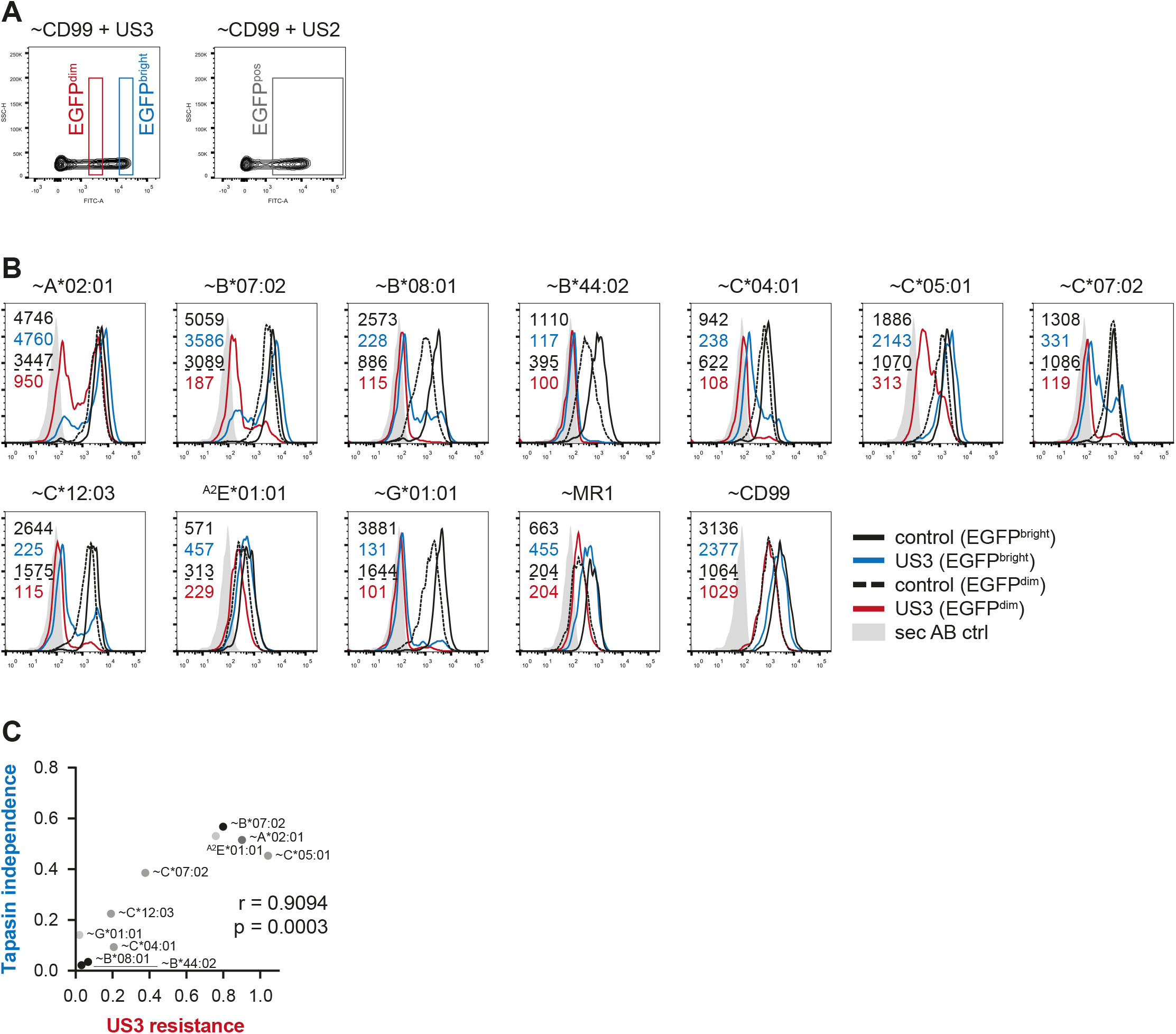
HLA-I regulation by US3 expressed in the EGFP^bright^ gate strongly correlates with HLA-I tapasin dependency. (A) EGFP-gating used for Fig. 5A-B (EGFP^pos^) and EGFP^dim^ and EGFP^bright^ cells in Fig. 5C. (B) Representative histograms for Fig. 5C, depicting distinct HLA-I expression in EGFP^dim^ and EGFP^bright^ gates. (C) Two-tailed correlation analysis of the results for EGFP^bright^ cells from Fig. 5C compared to the HLA-I tapasin independence analysis in Fig. 3A.

**Suppl. Fig. 5.**
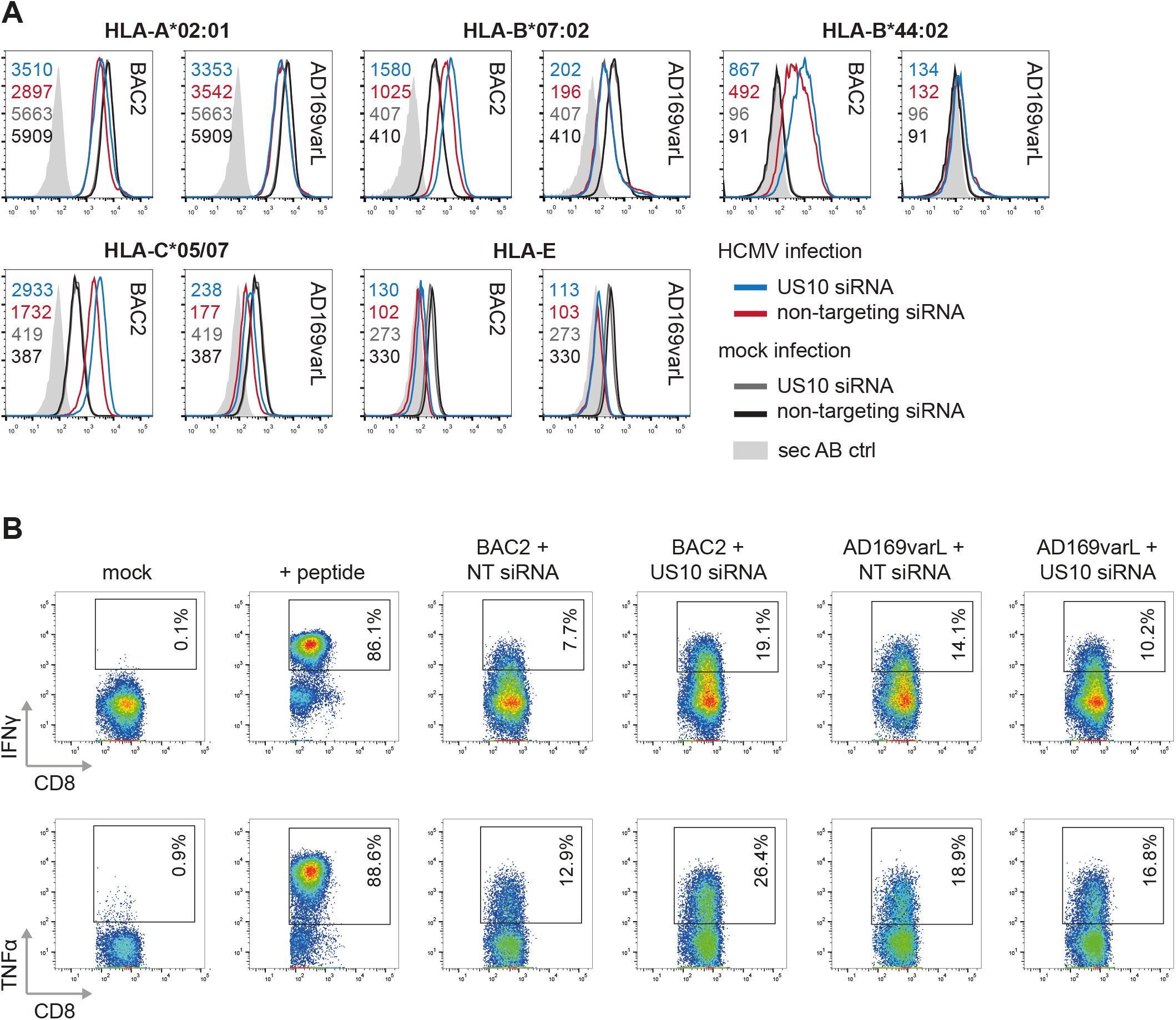
Slow turnover of US10; US10 knock-down by siRNA. (A) HeLa cells were transiently transfected with a US10-encoding plasmid or a control plasmid (-). At 24 h post-transfection, cells were metabolically labeled for 30 min and chased as indicated. Immunoprecipitations were performed with an anti-US10 antiserum and retrieved proteins were separated by SDS-PAGE and detected by autoradiography. EndoH was applied as indicated. One of two independent experiments (biological replicates) is shown. (B) Depiction of the coding sequences (CDS) for US11 and US10 together with the binding sites of the primers used for Fig. 6C and of US10-targeting siRNA #1. (C) Stable HeLa-US10HA cells were transfected with three different siRNA targeting US10 (#1-3) or with non-targeting (NT) siRNA. Whole cell lysates were prepared for Western blot analysis using anti-HA antibodies. US10-targeting siRNA #1 was used for all subsequent experiments. One of two independent experiments (biological replicates) is shown. (D-E) MRC-5 fibroblasts were nucleofected with US10HA, US3HA or a control together with US10- or non-targeting siRNA. At 24h post-nucleofection, an intracellular staining was performed with anti-HA and analyzed by flow cytometry. Fold change by US10 siRNA is depicted in (E). Dots represent individual values from two independent experiments (biological replicates).

**Suppl. Fig. 6.**
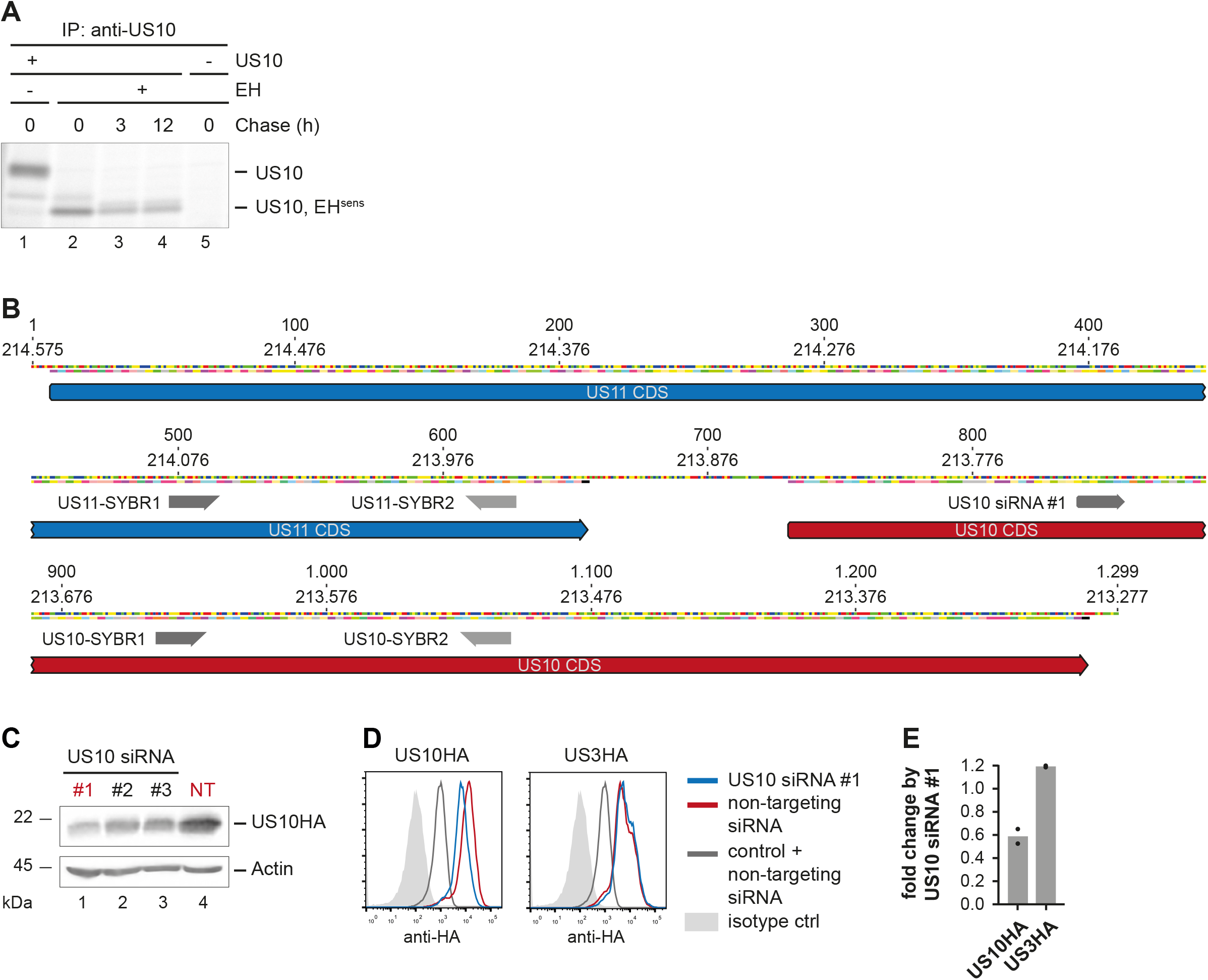
HLA-I regulation in US10 siRNA-treated HCMV-infected fibroblasts. (A) Representative histograms from Fig. 7A. (B) Representative dot plot for Fig. 7B. T-cells incubated with the pp65 peptide were used as positive control (+peptide).

